# VASH1-SVBP and VASH2-SVBP generate different detyrosination profiles on microtubules

**DOI:** 10.1101/2022.06.02.494516

**Authors:** Sacnicte Ramirez-Rios, Sung Ryul Choi, Chadni Sanyal, Thorsten Blum, Christophe Bosc, Fatma Krichen, Eric Denarier, Jean-Marc Soleilhac, Béatrice Blot, Carsten Janke, Virginie Stoppin-Mellet, Maria M. Magiera, Isabelle Arnal, Michel O. Steinmetz, Marie-Jo Moutin

**Affiliations:** Univ. Grenoble Alpes, Inserm, U1216, CNRS, CEA, Grenoble Institut Neurosciences, 38000 Grenoble, France; Laboratory of Biomolecular Research, Division of Biology and Chemistry, Paul Scherrer Institut, 5232 Villigen PSI, Switzerland; Institut Curie, Université PSL, CNRS UMR3348, 91401 Orsay, France; Université Paris-Saclay, CNRS UMR3348, 91401 Orsay, France; Biozentrum, University of Basel, 4056 Basel, Switzerland

**Keywords:** Microtubule, tubulin, detyrosination, tyrosination, vasohibin, VASH, SVBP

## Abstract

The detyrosination/tyrosination cycle of α-tubulin is critical for proper cell functioning. VASH1-SVBP and VASH2-SVBP are ubiquitous enzyme complexes involved in microtubule detyrosination. However, little is known about their mode of action. Here, we show in reconstituted systems and in cells that VASH1-SVBP and VASH2-SVBP drive global and local detyrosination of microtubules, respectively. We solved the cryo-electron microscopy structure of human VASH2-SVBP bound to microtubules, revealing a different microtubule-binding configuration of its central catalytic region compared to VASH1-SVBP. We further show that the divergent mode of detyrosination between the two enzymes is correlated with the microtubule-binding properties of their disordered N- and C-terminal regions. Specifically, the N-terminal region is responsible for a significantly longer residence time of VASH2-SVBP on microtubules compared to VASH1-SVBP. We suggest that this VASH domain is critical for microtubule-detachment and diffusion of VASH-SVBP enzymes on the lattice. Together, our results suggest a mechanism by which these enzymes could generate distinct microtubule subpopulations and confined areas of detyrosinated lattices to drive various microtubule-based cellular functions.

**SUMMARY:** VASH1-SVBP and VASH2-SVBP produce global and local detyrosination patterns of microtubule lattices, respectively. These activities rely on the interplay between the N- and C-terminal disordered regions of the enzymes, which determine their differential molecular mechanism of action.

**GRAPHICAL ABSTRACT:** Schematic representation of divergent molecular mechanisms of action of VASH-SVBP detyrosination complexes.

## INTRODUCTION

Microtubules are key eukaryotic cytoskeletal elements that are regulated by the tubulin code. This code involves the differential expression of α- and β-tubulin isotypes and their post-translational modifications (reviewed in (Janke and Magiera, 2020)). A prominent tubulin post-translational modification is the cyclic removal and re-addition of the C-terminal tyrosine residue of α-tubulin. This evolutionary conserved cycle, referred to as the “detyrosination/tyrosination cycle”, was shown to be crucial for cell mitosis and differentiation; it is also a key player in the development and functioning of the brain and the heart ((Chen et al., 2018; Erck et al., 2005), reviewed in (Lopes and Maiato, 2020; Moutin et al., 2021; Sanyal et al., 2021)). Accordingly, dysfunctions of the detyrosination/tyrosination cycle lead to cancer, brain disease, and heart failure in humans ((Chen et al., 2020; Lafanechere et al., 1998; Peris et al., 2022; Schuldt et al., 2021) reviewed in (Lopes and Maiato, 2020; Sanyal et al., 2021)). The known “readers” of the tubulin tyrosine signal emerging from the cycle are kinesin motor and CAP-Gly proteins (reviewed in (Sanyal et al., 2021; Steinmetz and Akhmanova, 2008). The activity of the detyrosination/tyrosination cycle regulates cargo trafficking in cells by tuning the microtubule binding of specific kinesins and the dynein-dynactin complex ((Dunn et al., 2008; Nirschl et al., 2016), reviewed in (Moutin et al., 2021)). The cycle also modulates tubulin interactions with regulators of microtubule stability, such as kinesin 13 and CLIP-170 (Chen et al., 2021; Peris et al., 2006; Peris et al., 2009). This modification is thus intimately linked to microtubule dynamics, where the detyrosination signal represents a marker of microtubule stability.

The detyrosination/tyrosination cycle of α-tubulin involves at least four enzymes including the tubulin tyrosine ligase (TTL, (Ersfeld et al., 1993)) and the three recently discovered detyrosinases comprising the two enzymatic complexes composed of a vasohibin (VASH1 or VASH2) and a small vasohibin-binding protein (SVBP) (Aillaud et al., 2017; Nieuwenhuis et al., 2017), and MATCAP (microtubule associated tubulin carboxypeptidase (Landskron et al., 2022)). TTL represents the “writer” of the tyrosine signal, while vasohibins and MATCAP are the “erasers”. SVBP acts as a chaperone and a co-factor for VASH1 and VASH2, which are specific tyrosine/phenylalanine carboxypeptidases (Aillaud et al., 2017; Nieuwenhuis et al., 2017). Mammalian VASH1 and VASH2 proteins share more than 50% overall sequence identity. They belong to the transglutaminase-like cysteine protease superfamily (Sanchez-Pulido and Ponting, 2016) and harbor a Cys–His–Leu catalytic amino acid residue triad conserved among species (Adamopoulos et al., 2019; Wang et al., 2019). Crystal structures of VASH1-SVBP and VASH2-SVBP complexes at high resolution allowed to determine the global organization of the central catalytic regions of the VASHs (also called core domains), which are their most homologous domains ((Adamopoulos et al., 2019; Li et al., 2019; Liao et al., 2019; Liu et al., 2019; Wang et al., 2019; Zhou et al., 2019), reviewed in (Sanyal et al., 2021). These domains form a folded, globular structure enclosing the C-terminal region of SVBP. The N- and C-terminal regions of VASHs are less well conserved and are assumed to be disordered as underlined by the lack of structural information in all published structures of VASH-SVBP complexes to date (Adamopoulos et al., 2019; Li et al., 2019; Liao et al., 2019; Liu et al., 2019; Wang et al., 2019; Zhou et al., 2019).

VASH-SVBP complexes are known to preferentially detyrosinate microtubule-lattice incorporated α-tubulin (Aillaud et al., 2017). We have previously shown that the VASH2-SVBP complex binds microtubules with a 1:1 stoichiometry of VASH2-SVBP:αβ-tubulin heterodimer and with an equilibrium dissociation constant in the low micromolar range (Wang et al., 2019). Mutating the catalytic cysteine residue Cys158 of VASH2 resulted in an enzymatically inactive VASH2 version that was unable to detyrosinate α-tubulin, yet retaining the ability to co-localize with microtubules *in vitro* and in cells. We further identified positively charged VASH2 surface residues located in close proximity to the tyrosine-binding groove. Mutating these VASH2 residues decreased the detyrosination activity of the enzyme but did not completely abrogate microtubule binding *in vitro* and in cells, leading us to suspect that additional contact sites between VASHs and microtubules remained to be identified. In a recent work, cryo-electron microscopy (cryo-EM) was used to analyze the structure of VASH1-SVBP bound to microtubules. The authors showed that the core domain of VASH1 forms specific contacts with the folded parts of α-tubulin (Li et al., 2020). However, as for the crystal structures obtained on the isolated, free enzyme complexes, they did not resolve the flanking regions of the VASH1 core domain, again demonstrating that the N- and C-terminal regions of VASHs are disordered and flexible.

VASH proteins are ubiquitously expressed but at different basal levels, transcripts of VASH1 being in general much higher represented (around 10-times) than transcripts of VASH2 (see the Human Protein Atlas project portal). In cells, detyrosination can occur on microtubules for many purposes and in several places (for review see (Roll-Mecak, 2019)), and the precise cellular role of each VASH-SVBP complex remains to be understood. VASH1-SVBP was identified as a primary tubulin detyrosinase in neurons and cardiomyocytes (Aillaud et al., 2017; Chen et al., 2020); however, VASH2-SVBP seems to act in cardiac cells as well (Yu et al., 2021). Thus, although a large amount of structural information has been obtained, the mechanism of action of VASH-SVBP enzyme complexes remain poorly understood. Here, combining single-molecule total internal reflection microscopy (TIRF) assays and cryo-EM, we depicted the molecular basis of the interaction of VASH1-SVBP and VASH2-SVBP with microtubules in relationship with their detyrosination activities. We showed that VASH1-SVBP generates global detyrosination of microtubule while VASH2-SVBP leads to local modifications both in vitro and in cells. The in vitro results obtained with several truncated mutants unveil crucial roles of the disordered N- and C-terminal regions in differentially regulating VASHs’ interactions with microtubules and hence their activities. We reveal that the N-terminal regions, which are highly divergent between the two VASHs, contribute to the difference in the enzymes’ ability to diffuse on the microtubule lattice. Our data further suggest that the disparity in functioning of the two VASH-SVBP enzymes contributes to the generation of different patterns of detyrosinated microtubules and thus to microtubule subpopulations having distinct roles within cells.

## RESULTS

### VASH1 and VASH2 drive global and local microtubule detyrosination, respectively

We first set up experimental conditions for which both the detyrosinating activity as well as the interaction with microtubules of VASH-SVBP complexes could be studied in parallel using immunofluorescence and TIRF microscopy, respectively. We generated constructs encoding the active full-length human VASH1 or VASH2 tagged with sfGFP at their N-termini and myc-6His at their C-termini. These constructs, together with the plasmid encoding the human SVBP fused to myc and FLAG tags at the C-terminus, were used to prepare full-length sfGFP-VASH1/2-SVBP complexes (V1_FL and V2_FL; Figure S1AB). We immobilized Taxol-stabilized microtubules enriched in tyrosinated HeLa tubulin (Tyr-MTs, containing 80 % of tyrosinated-tubulin, see Materials and Methods and Figure S1C) on the surface of TIRF chambers as previously described (Stoppin-Mellet et al., 2020). Different media were tested in order to find experimental conditions where single molecules of the two enzyme complexes associated with microtubules. In BRB40 (40 mM PIPES-KOH at pH 6.8, 1 mM MgCl_2_, 1 mM EGTA) supplemented with 50 mM KCl, both enzyme complexes were fully soluble, and showed activity and binding events at 50 pM protein concentration, allowing to carry out a comparative study.

We first tested the activity of VASH1-SVBP and VASH2-SVBP in vitro using immunofluorescence (Figure 1 A-C). Active versions of the two enzyme complexes (V1_FL and V2_FL) were incubated with Taxol-stabilized microtubules enriched in tyrosinated HeLa tubulin (Tyr-MTs) for 30 min. Both complexes led to a decrease of tyrosinated and a rise of detyrosinated microtubules (Figure 1A). Quantification of tyrosination and detyrosination levels showed that the changes were significantly larger for VASH1-SVBP than for VASH2-SVBP (Figure 1B). Interestingly, while the VASH1-SVBP complex induced a global microtubule detyrosination, VASH2-SVBP induced the formation of local areas of modifications of the microtubule lattice (Figure 1AC). Extensive regions of microtubules remained unmodified upon incubation with VASH2-SVBP over a 30 min period, which was not the case for VASH1-SVBP. In order to avoid a contribution of the fusion tags present in both the VASHs on their behavior, we tested enzymatic complexes with no tags on the VASHs and only a 6xHis tag at the N-terminus of SVBP. GFP-untagged proteins behaved similarly to their tagged counterparts: The VASH1-SVBP complex induced global detyrosination while VASH2-SVBP led to spots of detyrosination (Figure S2A-D).

**Figure 1.**
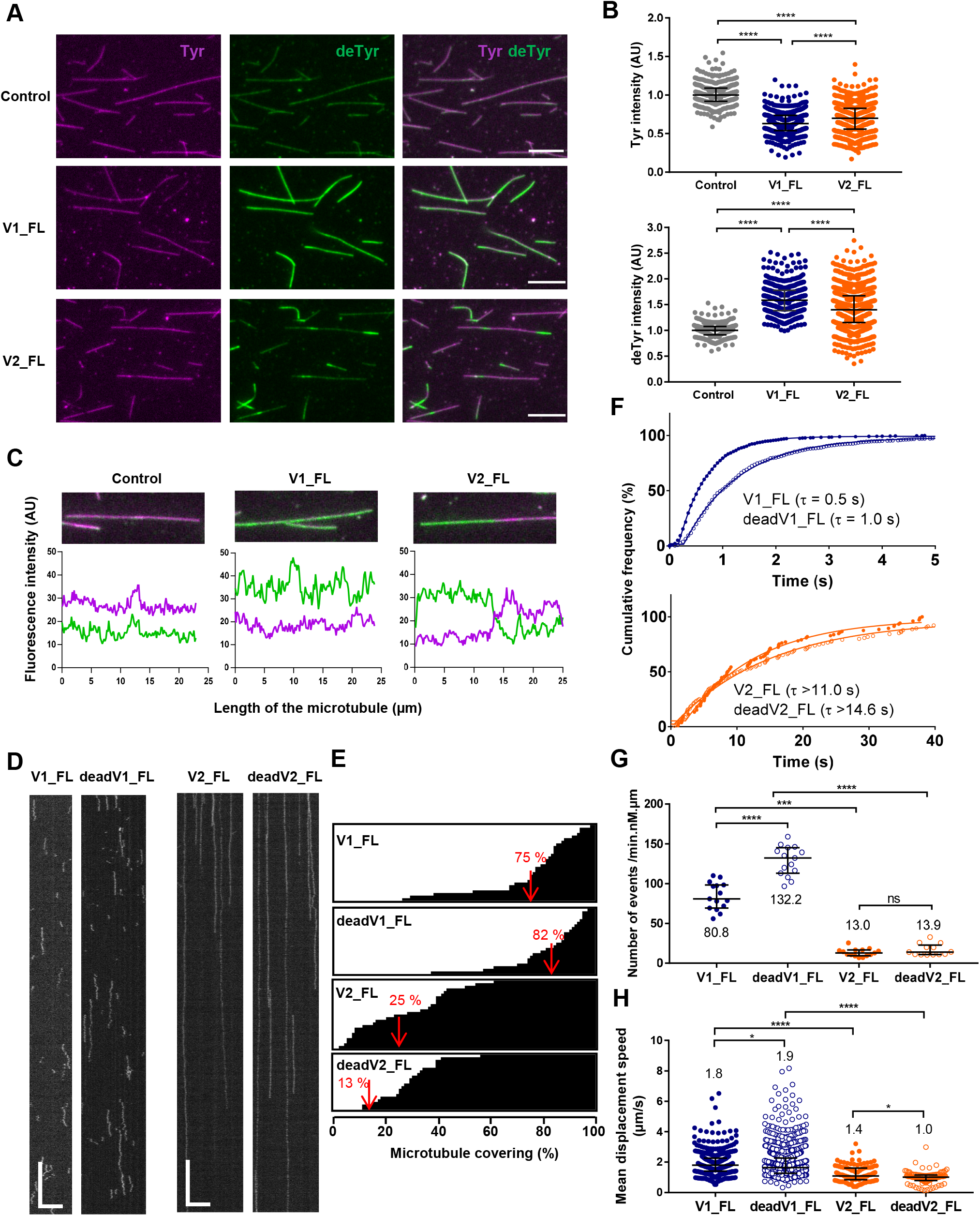
VASH1-SVBP and VASH2-SVBP complexes exhibit dissimilar microtubule detyrosination activities and very different binding behaviors on microtubules. **(A-C)** Comparison of the activity of sfGFP-tagged VASH1-SVBP (V1_FL) and VASH2-SVBP (V2_FL) enzyme complexes (50 pM) on Taxol-stabilized microtubules enriched in HeLa tyrosinated tubulin (Tyr-MTs) measured by immunofluorescence in BRB40 supplemented with 50 mM KCl. **(A)** Representative images of tyrosinated (Tyr, magenta) and detyrosinated (deTyr, green) α-tubulin pools of microtubules after 30 min incubation in the absence (control) or presence of the indicated enzyme complexes. Scale bar, 10 µm **(B)** Analysis of tyrosinated-and detyrosinated-tubulin signal intensity. Each point represents a microtubule (at least 300 microtubules were analyzed). Data are represented as the median with the interquartile range. Statistical significance was determined using Kruskal-Wallis test, ****p < 0.0001. **(C)** Graphs of fluorescence intensity variations of tyrosinated (magenta) and detyrosinated (green) tubulin on selected microtubules in the absence and presence of the enzyme. **(D-G)** TIRF microscopy study of single molecules of sfGFP-tagged VASH1-SVBP (V1_FL and deadV1_FL) and VASH2-SVBP (V2_FL and deadV2_FL) bound to microtubules in the same experimental conditions as in A-C. **(D)** Representative kymographs. Scale bars: horizontal, 5 µm; vertical, 5 s. Examples of TIRF movies for active VASH1/2-SVBP from which kymographs were extracted are presented in supplemental data (Videos S1, S2). (**E-G**) Analysis of the binding characteristics. Results for VASH1-SVBP are in blue and for VASH2-SVBP in orange, with plain circles for active enzymes (V1/2_FL) and empty circles for their catalytically dead versions (deadV1/2_FL). **(E)** Representation of the microtubule surface covered with VASH-SVBP molecules (white) or not covered (black) during the 45 s of a TIRF movie. Each horizontal line represents a microtubule (at least 19 microtubules were analyzed). The red value corresponds to the mean covering (in %). **(F)** Cumulative frequency of the residence times measured in TIRF movies taken during the 30 min following addition of enzyme complexes to microtubules. The mean residence time (τ) is obtained by fitting the curve with a mono-exponential function. **(G)** Analysis of binding frequency. Each point represents a microtubule. **(H)** Analysis of diffusion. Each point represents a single molecule, the molecules moving on at least 15 microtubules were analyzed. Data are represented as the median with the interquartile range. Statistical significance was determined using Kruskal-Wallis test. *p < 0.01, ***p < 0.005, ****p < 0.0001, and ns, not significant. Binding characteristics are also summarized in Table 1.

Together, these results show that VASH1-SVBP or VASH2-SVBP effectively detyrosinates microtubules in vitro, however, in a different manner.

### Difference in the interaction of VASH1-SVBP and VASH2-SVBP with microtubules

In order to understand the molecular mechanisms involved in the distinct detyrosination patterns, we monitored the interaction of the full-length VASH1-SVBP and VASH2-SVBP enzyme complexes (V1_FL and V2_FL) with Taxol-stabilized microtubules enriched in tyrosinated HeLa tubulin (Tyr-MTs) in BRB40 supplemented with 50 mM KCL. We also examined the microtubule-interacting behavior of catalytically inactive enzyme complexes containing a Cys169Ala mutation in VASH1 (deadV1_FL) or a Cys158Ala mutation in VASH2 (deadV2_FL) (Aillaud et al., 2017; Wang et al., 2019). Interestingly, as shown in representative kymographs, the binding behavior of the two enzyme complexes on microtubules was very different (Figure 1D). Whereas both active and dead versions of VASH1-SVBP exhibited short and frequent binding events, active and dead versions of VASH2-SVBP bound less frequently and for much longer times on microtubules (Videos S1 and S2). As illustrated in Figure 1E, the local activity of VASH2-SVBP was correlated with poor covering of microtubules by this enzyme complex in TIRF experiments carried out under the same conditions. In contrast, microtubule covering by VASH1-SVBP enzymes was high (75% and 82% covering for V1_FL and deadV1_FL, respectively, versus 25% and 13 % covering for V2_FL and deadV2_FL, respectively).

We next analyzed the binding parameters (residence-time, binding frequency, diffusion) of the enzyme complexes on microtubule lattices in these experimental conditions (Figure 1F-H and Table 1). Within 30 min after the addition of the enzyme complex, the residence-time distribution of VASH1-SVBP on microtubules was monoexponential with a mean residence-time (τ) of 0.5 and 1.0 s for the active and dead enzyme versions, respectively. The mean residence-time of full-length VASH2-SVBP was estimated higher than 11 s for the active version and higher than 14.6 s for the dead version, and thus at least 15- to 22-times higher than the residence-time of VASH1-SVBP (Figure 1F). A precise determination of the residence time for VASH2-SVBP was not possible in the experimental buffer conditions used to study both complexes side by side. Indeed, most VASH2-SVBP molecules were already attached to microtubules at the beginning of the TIRF movies and single molecule traces disappeared during movie acquisition due to quenching of the sfGFP fluorophore (see Materials and Methods for details). As illustrated in Figure 1G, VASH2-SVBP displayed a 6- to 10-times lower binding frequency than VASH1-SVBP (13.0 events/min.µm.nM of V2-FL and 13.9 events/min.µm.nM of deadV2-FL to be compared to 80.8 events/min.µm.nM of V1_FL and 132.2 events/min.µm.nM of deadV1_FL). Both enzymes diffused along the microtubule lattice, with VASH2-SVBP diffusing more slowly than VASH1-SVBP (1.8 and 1.9 µm.sec^-1^ for V1_FL and deadV1_FL, respectively to be compared to 1.4 and 1.0 µm.sec^-1^ for V2_FL and deadV2_FL, respectively, Figure 1H).

**Table 1.** Binding parameters of VASH-SVBP complexes interaction with tyrosinated- and detyrosinated-enriched microtubules in variable ionic strength media. **(A)** Related to Figure 1D-G. **(B)** Related to Figure 2A. **(C)** related to Figure 2B. **(D)** related to Figure 2C and 3CD. **(E)** related to Figures 3AB. A, B, C, D, E represent experiments performed on different days. At least 15 microtubules were analyzed. See Materials and Methods for detailed experimental conditions.

| Experiment (medium) | Ionic strength (mM) | Microtubule | Recombinant protein (concentration) | Residence time $\tau$ (s) | Binding frequency ($\text{min}^{-1} \cdot \text{nM}^{-1} \cdot \mu\text{m}^{-1}$) |
| --- | --- | --- | --- | --- | --- |
| **A** (BRB40, 50 mM KCl) | 140 | Tyr-MTs | V1_FL (50 pM) | 0.5 | 80.8 |
|  |  | Tyr-MTs | deadV1_FL (50 pM) | 1.0 | 132.2 |
|  |  | Tyr-MTs | V2_FL (50 pM) | >11 | 13.0 |
|  |  | Tyr-MTs | deadV2_FL (50 pM) | >14.6 | 13.9 |
| **B** (BRB80) | 173 | Tyr-MTs | V1_FL (50 pM) | 0.8 | 51.6 |
|  |  | Tyr-MTs | deadV1_FL (50 pM) | 0.9 | 86.4 |
| **C** (BRB40, 50 mM KCl) | 223 | Tyr-MTs | V1_FL (1 nM) | 0.2 | 3.0 |
|  |  | Tyr-MTs | deadV1_FL (1 nM) | 0.2 | 12.7 |
|  |  | Tyr-MTs | V2_FL (100 pM) | 3.9 | 20.4 |
|  |  | Tyr-MTs | deadV2_FL (100 pM) | >7.2 | 13.5 |
| **D** (BRB40, 50 mM KCl) | 273 | Tyr-MTs | V2_FL (50 pM) | 0.9 | 14.9 |
|  |  | Tyr-MTs | deadV2_FL (50 pM) | 1.4 | 19.3 |
|  |  | deTyr-MTs | V2_FL (50 pM) | 0.6 | 13.7 |
|  |  | deTyr-MTs | deadV2_FL (50 pM) | 0.7 | 41.0 |
| **E** (BRB40, 50 mM KCl) | 140 | Tyr-MTs | V1_FL (50 pM) | 0.8 | 53.4 |
|  |  | Tyr-MTs | deadV1_FL (50 pM) | 1.2 | 89.0 |
|  |  | deTyr-MTs | V1_FL (50 pM) | 0.5 | 42.3 |
|  |  | deTyr-MTs | deadV1_FL (50 pM) | 0.6 | 66.5 |

As the binding of microtubule-associated proteins to microtubules often involves electrostatic attractive interactions, we tested whether an increase in ionic strength affects the interaction of VASH-SVBP with microtubules. This significantly decreased the run length of both enzyme complexes on microtubules (Figure 2A-C and Table 1), and allowed to determine the residence-time for VASH2-SVBP binding (Figure 2C and Table 1D). Because it does not alter the evaluation of residence time and allows better visibility of traces on microtubules, the concentration of VASH1-SVBP was increased when using high ionic strength buffers. Despite reducing the residence time, VASH2-SVBP still stayed much longer on microtubules than VASH1-SVBP (Table 1C, τ = 0.2 s for V1_FL versus τ = 3.9 s for V2_FL in BRB80 supplemented with 50 mM KCl, which represents more than 80 mM increase in ionic strength compared to BRB40 supplemented with 50 mM KCl). We next assayed the impact of ionic strength on the enzymatic activity by immunofluorescence. Quantifications of the tyrosinated tubulin signal of images from experiments performed in varying buffers demonstrated that, like the binding behavior, the enzymatic activity of the two VASH-SVBP complexes was salt dependent: increasing the ionic strength led to a significant reduction of the enzymatic activity (Figure 2D).

**Figure 2.**
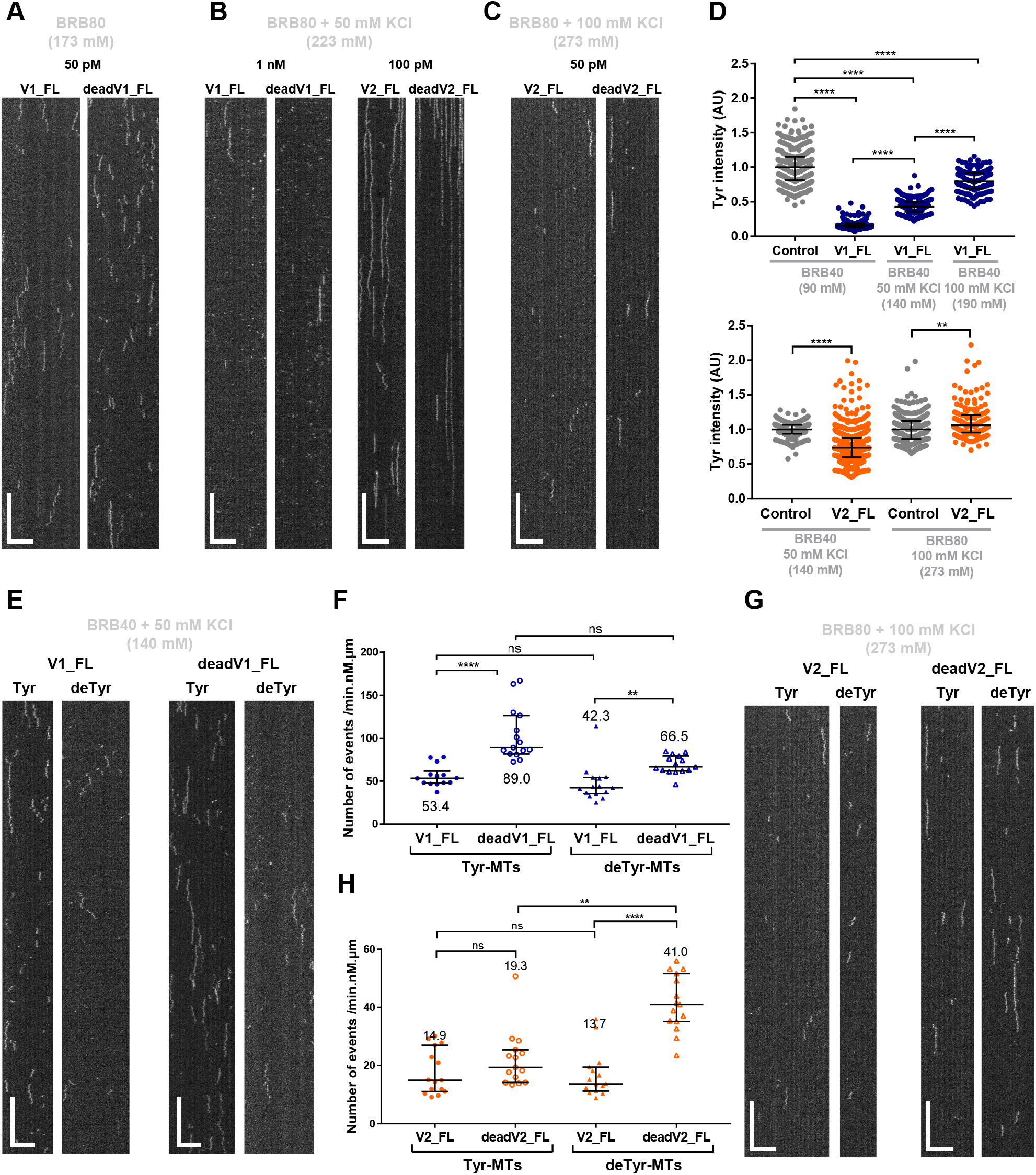
Ionic strength has a strong impact on the interaction of VASH-SVBP complexes with microtubules, whereas tyrosinated-tubulin depletion disrupts less the interaction. **(A-D)** Effect of ionic strength. Experiments were performed in the indicated buffers. Ionic strengths are indicated in brackets in mM. **(A-C)** TIRF microscopy study. Representative kymographs of single molecules of sfGFP-tagged VASH1-SVBP and VASH2-SVBP (V1_FL, deadV1_FL, V2_FL and deadV2_FL) at the indicated concentrations bound to Taxol-stabilized microtubules enriched in HeLa tyrosinated tubulin (Tyr-MTs). Scale bars: horizontal, 5 µm; vertical, 5 s. Binding characteristics are summarized in Table 1BCD. **(D)** Activity measurement by immunofluorescence. Analysis of tyrosinated signals intensities after 30 min incubation in the absence (control) or presence of 50 pM enzymes. Each point represents a microtubule (at least 150 microtubules were analyzed). Data are represented as the median with the interquartile range. **(E-H)** Effect of decreasing tyrosinated tubulin. TIRF microscopy study of single molecules of sfGFP-tagged VASH1-SVBP (V1_FL and deadV1_FL, 50 pM) and VASH2-SVBP (V2_FL and deadV2_FL, 50 pM) bound to Taxol-stabilized microtubules enriched in HeLa tyrosinated tubulin (Tyr-MTs) or detyrosinated tubulin (deTyr-MTs). Binding characteristics are summarized in Table 1. **(E, G)** Representative kymographs. Scale bars: horizontal, 5 µm; vertical, 5 s. **(F, H)** Analysis of binding frequency. Each point represents a microtubule. At least 15 microtubules were analyzed. Data are represented as the median with the interquartile range. Statistical significance was determined using Kruskal-Wallis test. *p < 0.01, **p < 0.009, ***p < 0.005, ****p < 0.0001, and ns, not significant.

Together, these results clearly highlight the different binding behaviors of the two VASH-SVBP enzyme complexes and the importance of electrostatic attractive interactions for their microtubule binding.

### Tyrosination has a little effect on the interaction of VASH-SVBP complexes with microtubules

We then wondered how VASH-SVBP complexes behave on detyrosinated, non-substrate microtubules. We compared the interaction of both enzyme complexes with microtubules enriched either in tyrosinated or detyrosinated HeLa tubulin (Tyr-MTs contained 80 % tyrosinated tubulin, 17 % detyrosinated tubulin, and 3 % Δ2-tubulin (a variant missing the two C-terminal glutamate residues); deTyr-MTs contained 82 % detyrosinated tubulin, 15 % tyrosinated tubulin, and 3 % Δ2-tubulin; Figure S1C and Materials and Methods). We tested both active and inactive enzymes to assay the relevance of detyrosination activity on microtubule binding. To obtain complete traces and determine the binding characteristics of the two enzymes, a low ionic strength buffer (140 mM) was used for VASH1-SVBP and a high ionic strength buffer (273 mM) for VASH2-SVBP. Representative kymographs and binding characteristics are presented in Figure 2E-H and Table 1.

Catalytically active and dead versions of each enzyme complex displayed similar short residence times on deTyr-MTs (τ = 0.5 s for V1_FL and τ = 0.6 s for deadV1_FL in BRB40 supplemented with 50 mM KCl; τ = 0.6 s for V2_FL and τ = 0.7 s deadV2_FL in BRB80 supplemented with 100 mM KCl). Both enzyme versions stayed longer on Tyr-MTs than on deTyr-MTs, and dead enzyme complexes spent slightly longer times on Tyr-MTs than their active counterparts (τ = 0.8 s for V1_FL and τ = 1.2 s for deadV1_FL in BRB40 supplemented with 50 mM KCl; τ = 0.9 s for V2_FL and τ = 1.4 s for deadV2_FL in BRB80 supplemented with 100 mM KCl). The binding frequency of the catalytically dead VASH1-SVBP mutant was significantly higher than the one of its wild type counterpart on both Tyr-MTs and deTyr-MTs (Figure 2F). In contrast, the two VASH2-SVBP versions bound Tyr-MTs with similar frequencies (Figure 2H), which were 3 to 4-times lower than the ones of the two VASH1-SVBP versions. Surprisingly, the binding frequencies of catalytically dead VASH-SVBP mutants were higher on deTyr-MTs than on Tyr-MTs (Figure 2GH).

Thus, our results reveal slightly longer binding periods of both VASH1-SVBP and VASH2-SVBP complexes on tyrosinated microtubules than on detyrosinated ones (Table 1DE). This was best observed with the catalytically dead mutants that do not modify microtubules (two-fold longer binding). With active enzyme complexes, tyrosinated microtubules are progressively detyrosinated which, during the 30 min time of movie acquisition, could progressively reduce the residence time of the enzyme complexes (as shown for the highly efficient VASH1-SVBP enzyme complex; Figure S2EF). In the case of the catalytically dead VASH-SVBP versions, microtubules are not modified over the entire time of the experiment, allowing a detailed analysis of the interaction between enzymes and microtubules.

In addition, we found that at an appropriate ionic strength the two dead mutants were able to dissociate from the microtubule lattice, implying that the enzymes’ detyrosination activity is not essential for the release of these enzymes from microtubules. Notably, in such conditions catalytically active and dead versions of the two enzyme complexes showed diffusion in both directions along Taxol-stabilized, tyrosinated tubulin-enriched microtubules.

Collectively, these results suggest that in vitro, VASH1-SVBP exhibits short and diffusive binding events on microtubules while VASH2-SVBP displays longer and much more static attachments to them. In addition, the catalytic activity of the two enzyme complexes does not appear to be essential for their binding behavior on microtubules.

### Cryo-EM structure of VASH2-SVBP in complex with a microtubule

The functional results collected so far suggest that VASH1-SVBP and VASH2-SVBP work very differently on microtubules. We hypothesized that this could be related to a different binding mode of the two enzyme complexes with microtubules. To investigate this idea, we analyzed the structure of VASH2-SVBP in complex with microtubules by cryo-EM. We used the catalytically dead mutant of VASH2-SVBP and GMPCPP-stabilized HeLa cell microtubules (Figures 3A and S3). Based on a single particle approach (see Materials and Methods), we found that the majority of microtubules contained 14-protofilaments. Downstream extraction and single-particle classification of protofilaments revealed that VASH2-SVBP molecules were present in ∼40% of the total binding sites. Subsequent refinement of the particle subset generated an electron density map of two laterally associated αβ-tubulin heterodimers in complex with VASH2-SVBP at an overall resolution of 3.2 Å (Figure 3B; Figure S3B).

**Figure 3.**
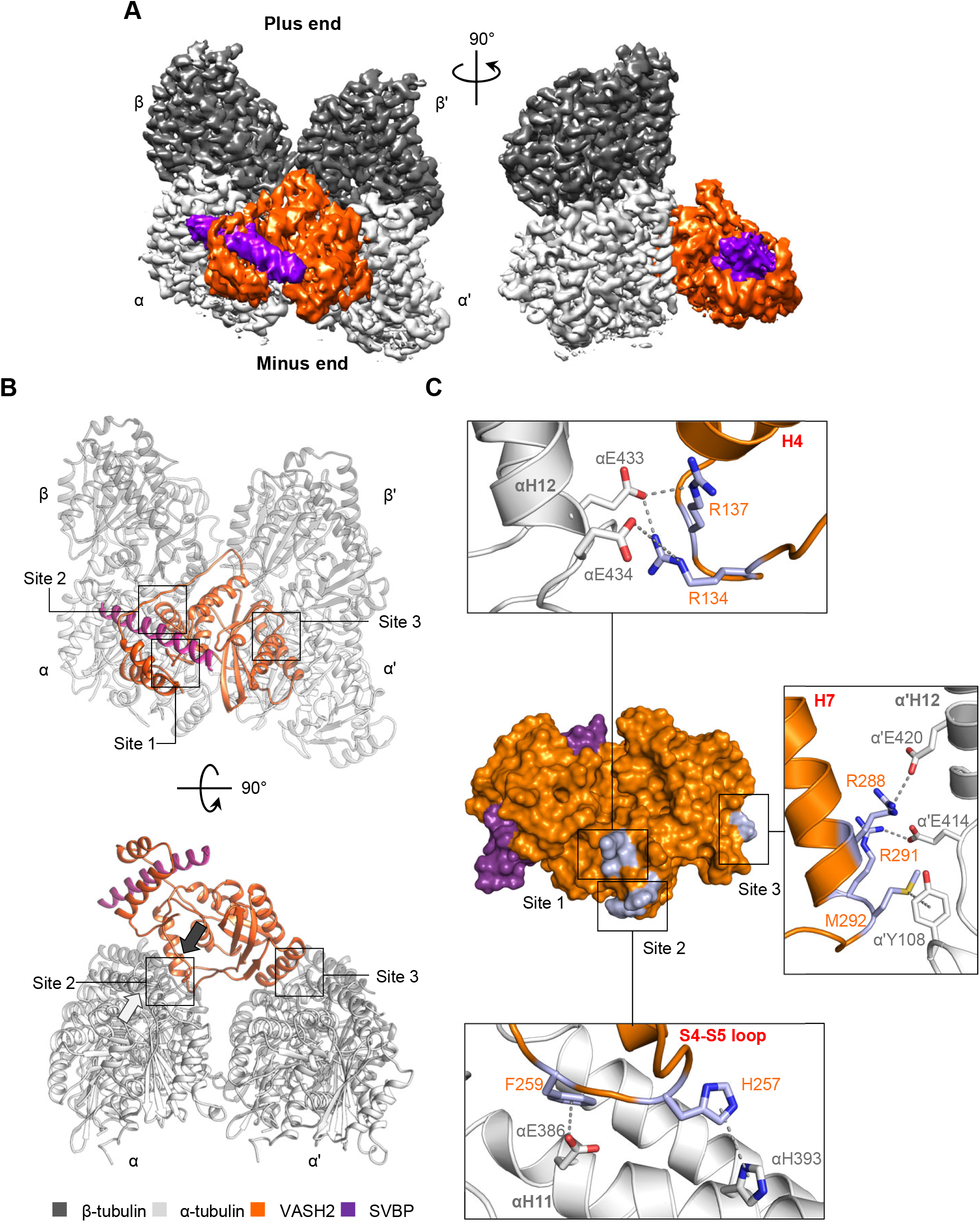
Cryo-EM reconstruction of the microtubule-VASH2-SVBP complex reveals their interaction sites. **(A)** Electron density map of VASH2-SVBP (catalytically dead version of VASH2 carrying the Cys158Ala mutation) bound to a GMPCPP-stabilized microtubule. A map region of two laterally associated αβ-tubulin dimers from two adjacent protofilaments bound to one VASH2-SVBP complex is shown in two different orientations 90° apart. **(B)** Overview of the three microtubule-VASH2-SVBP interacting sites from two different orientations 90° apart. Sites 1 and 2 involve the substrate α-tubulin while site 3 involves the α’-tubulin subunit of an adjacent protofilament in the microtubule. All molecules are displayed in cartoon representation. The white arrow close to site 2 in the lower panel indicates the C-terminus of helix αH12 and the black arrow denotes the facing direction of the positively charged groove on VASH2. **(C)** Surface representation of the VASH2-SVBP structure with the microtubule-binding residues highlighted in orange stick. Insets correspond to VASH2 and tubulin residues implicated in the three interaction sites, displayed in stick representation and labeled in orange and gray, respectively. The α-tubulin, β-tubulin, VASH2, and SVBP molecules are shown in light gray, dark gray, orange, and violet, respectively.

In the final map, we could reliably distinguish the α- and β-tubulin subunits by examining their corresponding S9-S10 loop densities (Figure S3D), a secondary structure element that differs in length between both subunits. The resolution of VASH2-SVBP was in the range of 3.3-4.8 Å becoming progressively higher in regions that are located closer to the microtubule surface (Figure S3C). It was straightforward to model unambiguously all regular secondary structure elements of all four protein components, α-tubulin, β-tubulin, VASH2, and SVBP, into the final electron density map, as well as a large number of residue side chains and both the tubulin guanosine nucleotides (Figure S3DE). Due to their intrinsic flexibility, the densities of the α- and β-tubulin C-terminal tails were defined poorly beyond the corresponding H12 helices and could thus not be modelled; the same also applied for the N- and C-termini of VASH2 and N-terminus of SVBP. The cryo-EM structure of the microtubule-bound VASH2-SVBP is very similar to the one of the free enzyme solved by X-ray crystallography (Wang et al., 2019) (RMSD over 256 Cα atoms of 0.79 Å), suggesting that the core of the enzyme does not undergo global structural rearrangement upon binding. Notably, the observed protofilament-straddling interaction between VASH2-SVBP and the microtubule (Figure 3AB) is not feasible for the free αβ- tubulin heterodimer. Therefore, the characteristic binding mode of VASH2-SVBP with its microtubule-incorporated α-tubulin substrate underlies the mechanism behind the preferential activity of VASH2-SVBP for microtubules compared to free tubulin (Kumar and Flavin, 1981). Table 2 provides a summary of the MolProbity validation statistics for the final atomic model.

**Table 2.**
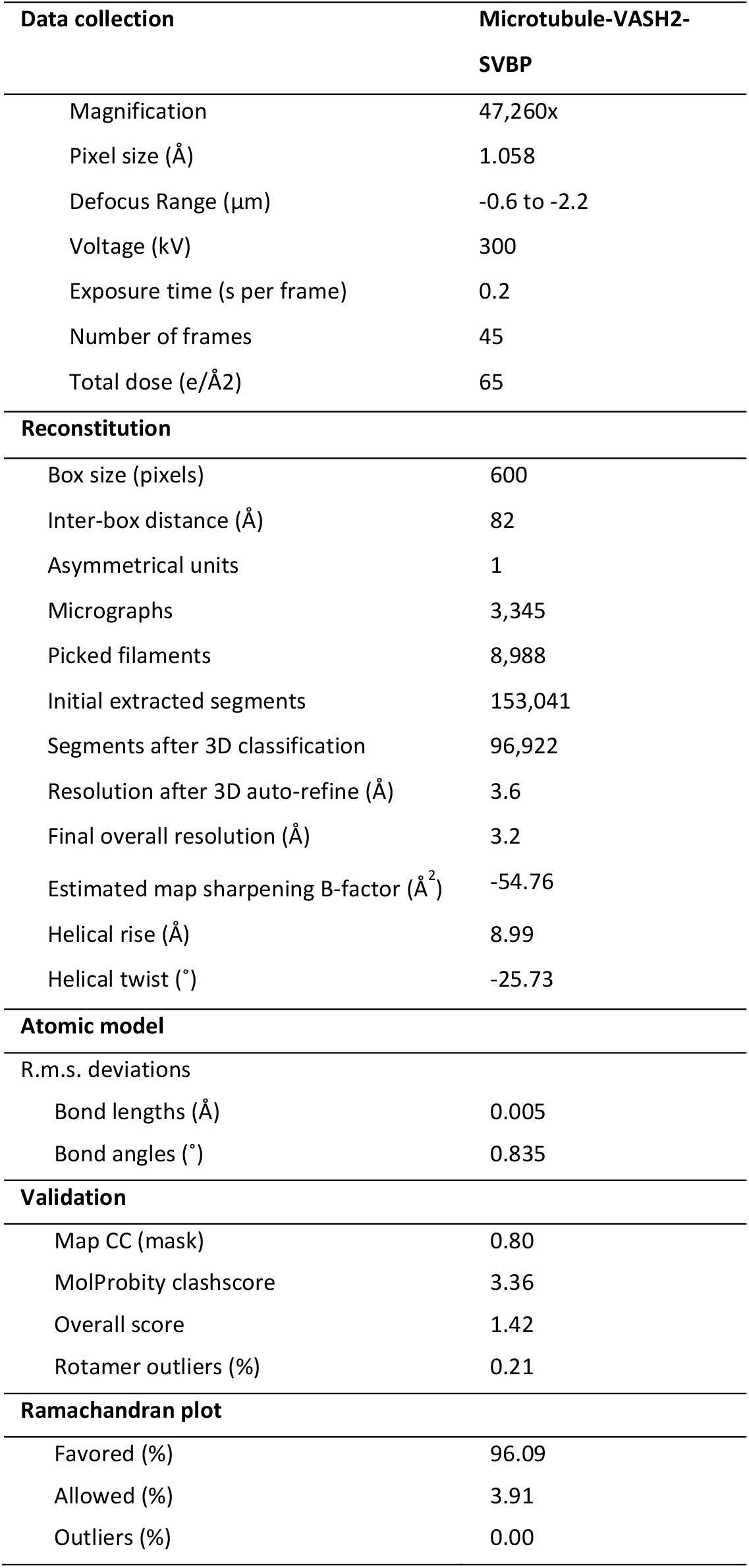
Cryo-EM data collection, refinement, and structure validation.

The model of the microtubule-VASH2-SVBP complex reveals a specific binding of the enzyme complex on the external surface of the microtubule. The central, positively charged groove of VASH2, encompassing the catalytic amino acid residue triad Cys158, His193 and Ser210 (Aillaud et al., 2017), faces towards the microtubule outer surface and is situated directly above the H12 helix of α-tubulin from which the flexible C-terminal tail emanates (not visible in the electron density; Figure 3B). Sites of contacts between VASH2-SVBP and the microtubule outer surface were located primarily on α-tubulin suggestive of specific recognition of the target substrate, the C-terminal tyrosine of α-tubulin. Conversely, the density corresponding to SVBP was distal from the microtubule-VASH2 binding interface and did not appear to contribute to the interaction. Interestingly, the VASH2 part of the enzyme complex binds simultaneously to two α-tubulin subunits: a α-tubulin subunit in close proximity to the VASH2 catalytic site and a α-tubulin subunit on the adjacent protofilament (henceforth denoted α’-tubulin; Figure 3B).

In support of our previous mutagenesis results (Wang et al., 2019), inspection of the microtubule-VASH2-SVBP complex structure revealed three primary sites of interaction, denoted site 1, site 2, and site 3, between the folded region of VASH2-SVBP and the ones of α- and α’-tubulin (Figure 3C). At site 1, residues Arg134 and Arg137 of VASH2 form electrostatic interactions with the negatively charged residues Glu434 and Glu433 of the H12 helix of α-tubulin, respectively. The S4-S5 loop of VASH2 is implicated in site 2 involving residue His257 and Phe259, which form polar interactions with His393 and Glu386 of α-tubulin, respectively. Site 3 engages the α’-tubulin subunit from the adjacent protofilament where a hydrophobic-aromatic residue interaction between Met292 of VASH2 and Tyr108 of α’-tubulin is implicated. In addition, residues Arg288 and Arg291 of VASH2 are likely to form electrostatic interactions with the negatively charged glutamate residues Glu420 and Glu414 of helix H12 of the α’-tubulin subunit, respectively.

Taken together, these results show that the structured domain of VASH2-SVBP is capable of binding to three distinctive sites on the microtubule surface. When bound to the microtubule, VASH2-SVBP interacts simultaneously with two adjacent protofilaments by binding both the α-tubulin substrate and the α’-tubulin of a neighboring protofilament. This interaction is driven by three binding sites that help position the positively charged groove of VASH2 in proximity to the C-terminal tail of α-tubulin. As the residues directly involved in microtubule-lattice binding are distinct and distal from the catalytic site, we may consider these as substrate recognition elements that confer specificity to the catalytic activity of the vasohibin family of tubulin carboxypeptidases.

### Comparison of the microtubule-binding mode of VASH1-SVBP and VASH2-SVBP

A cryo-EM structure of microtubule-bound VASH1-SVBP was recently solved based on a truncated, catalytically inactive version of VASH1 (Cys169 mutated to serine) and recombinant human tubulin (Li et al., 2020) (PDB ID 6WSL). Superimposition of the microtubule-bound structures of VASH2-SVBP and VASH1-SVBP revealed minimal structural differences between the two enzyme complexes (RMSD over 244 Cα atoms of 1.19 Å). As shown in Figure 4A, comparison of the microtubule-binding modes revealed that both enzyme complexes contact two adjacent α-tubulins across neighboring protofilaments simultaneously and position their catalytic site directly above the H12 helix of one of the two α-tubulin subunits. However, they do so at a different relative angle of approximately 24° as calculated from a plane defined by the Cα atoms of the three residues Val81, Glu148, and Glu284 of VASH2, and Ala92, Glu159, and Glu295 of VASH1.

**Figure 4.**
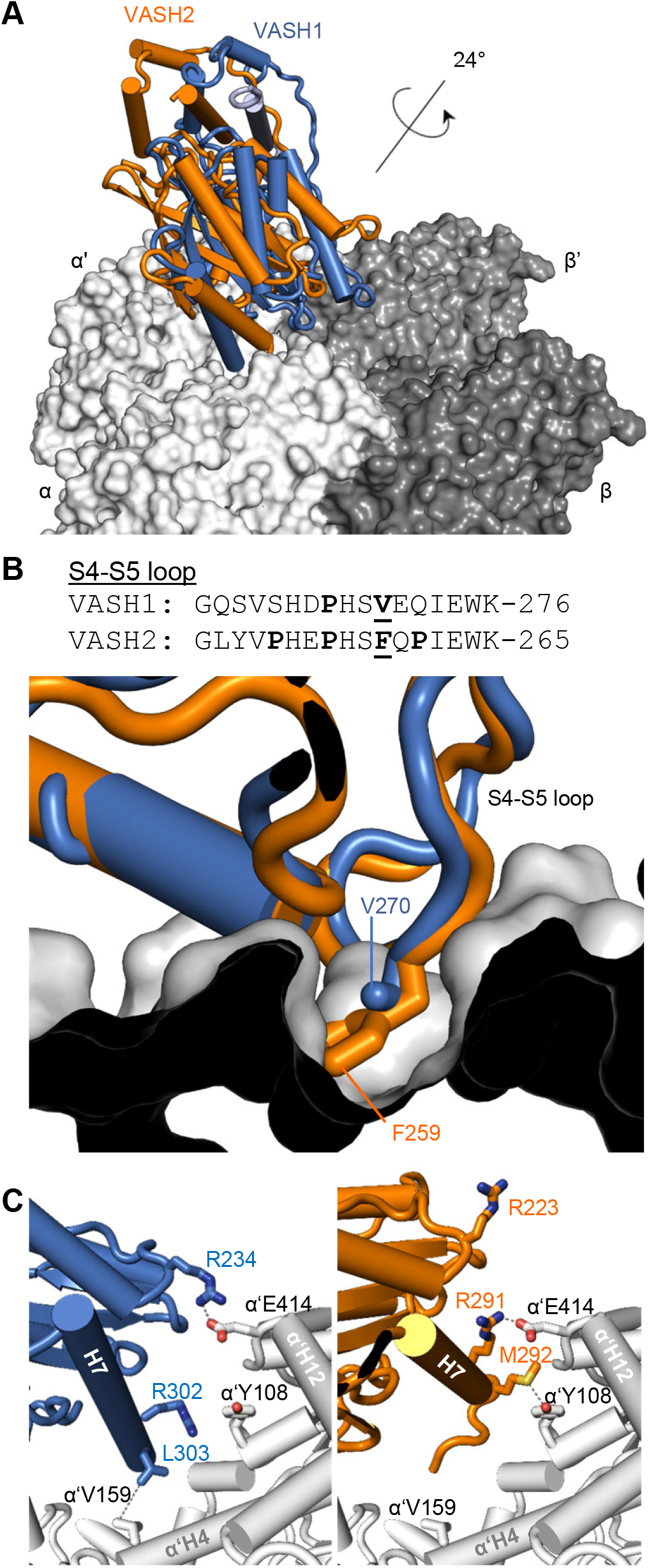
Comparison between the microtubule-binding modes of VASH1-SVBP and VASH2-SVBP. **(A)** Superimposition of the microtubule-VASH1-SVBP (PDB ID 6WSL, blue) and microtubule-VASH2-SVBP (orange) structures reveals a 24° tilt between the two enzyme complexes. **(B)** Superimposing the VASH2-SVBP structure (orange) onto the microtubule-bound VASH1-SVBP (blue) structure shows a steric clash of Phe259 of the β4-β5 loop of VASH2-SVBP with α-tubulin. On the top, a sequence alignment of the β4-β5 loops of VASH1 and VASH2 is shown with proline residues highlighted in bold. Phe259 of VASH2 and the corresponding Val270 of VASH1 are also highlighted in bold and are underlined. **(C)** At site 3, VASH1-SVBP (blue) residues Arg234 and Leu303 form contacts with residues Glu414 and Val159 of α’-tubulin, respectively. In the case of VASH2-SVBP (orange), residues Arg291 and Met292 engage residues Glu414 and Tyr 108 of α’-tubulin, respectively.

Although the sequence identity between full length VASH1 and VASH2 is high (58%, BLASTp analysis), the alignment of their sequences reveals non-identical residues at sites 2 and 3 (Figure S4), which may explain the observed difference in the overall microtubule-binding mode. Indeed, superimposition of the microtubule-bound structure of VASH2-SVBP onto that of VASH1-SVBP revealed that in the S4-S5 loop of VASH2, residue Phe259 (Val270 in VASH1) experiences a steric clash with His209 of the H9-S8 loop of α-tubulin (Figure 4B). Inspection of the amino acid sequence of the S4-S5 loop of VASH2 revealed the presence of two additional proline residues (Pro253 and Pro261) that are substituted by a serine and a glutamine, respectively, in VASH1 (Ser264 and Gln272; Figure 4B). We think that the additional rigidity imposed by these proline residues might force VASH2-SVBP to adapt a tilted microtubule-binding mode of 24° compared to VASH1-SVBP to accommodate the S4-S5 loop of VASH2. Binding site 3 displays additional differences between the structures of VASH2-SVBP and VASH1-SVBP, which may also account for the changes in the microtubule-binding mode of the two enzyme complexes. The methionine-aromatic interaction of Met292 of VASH2 with Tyr108 of α’-tubulin is not possible in VASH1 as the corresponding residue in VASH1 is a leucine (Leu303; Figure 4C). Furthermore, VASH2 does not engage the H4 helix of the α’-tubulin subunit as seen in the VASH1-SVBP structure (Figure 4C and (Wang et al., 2019)). Finally, the Arg234 residue in VASH1 is implicated in an electrostatic attractive interaction with Glu414 of the H12 helix of α’-tubulin; the corresponding residue in VASH2 (Arg223) is too distal from the α’-tubulin subunit to establish such an interaction (Figure 4C). Instead, Arg291 of VASH2 appears to be in close proximity to Glu414 of α’-tubulin and likely forms an electrostatic attractive interaction.

To determine what effect the conformational difference in microtubule binding may have on the enzymatic activity of the two VASH isoforms, we compared the relative position of the VASH1 and VASH2 catalytic sites (Figure S5). Using the coordinates of the catalytic triad, we triangulated the centroid point of the three Cα atoms of the corresponding residues as a representative position of the substrate. Compared to VASH2, the centroid point of VASH1 shifts by 6.3 Å in the direction of the plus-end and parallel to the protofilament. This places the catalytic cysteine of VASH1 at a distance of 29.7 Å from the Ser439 residue of the H12 helix of α-tubulin, which is comparable to the distance of 27.3 Å for VASH2. Residue Ser439 was selected, as it is the closest resolved residue of the α-tubulin C-terminus to the processed tyrosine. This analysis suggests that the observed difference in the microtubule-binding mode between VASH1 and VASH2 core domains has only minimal influence in the positioning of the catalytic site relative to the H12 helix of α-tubulin, and that the accessibility of the substrate C-terminal tail is thus only marginally changed between the two vasohibin isoforms.

Taken together, these results show that despite the similarities in the structures of the VASH1-SVBP and VASH2-SVBP core domains, a clear difference in their microtubule-binding pose is observed. These alternate poses are likely driven by the changes in the amino acid sequence of the S4-S5 loop between both enzymes, which forms interactions with site 2 of α-tubulin. Additional contribution may arise from the differences observed in the microtubule-binding residues at site 3 of α’-tubulin. However, this divergence could have only limited effect on the enzyme’s activity since the accessibility of the α-tubulin C-terminal tyrosine is quite similar.

### The core domains of VASH1 and VASH2 in complex with SVBP behave similarly

As the cryo-EM structures of microtubules in complex with VASH1-SVBP and VASH2-SVBP revealed specific interactions of the enzymatic core domains of VASHs with microtubules, we wonder how truncated versions of the enzymes containing only these central regions behave on microtubules in vitro. We thus examined the detyrosinating activity of VASH1 and VASH2 core domains in complex with SVBP (V1_CD and V2_CD, Figures 5AB and S1A), and used their catalytically dead versions (deadV1_CD and deadV2_CD) in order to avoid evolution of substrate over time to study their interaction with microtubules in single molecule TIRF experiments (Figure 5DE). A schematic representation of the two VASH proteins with their different domains is presented in Figure 5C.

**Figure 5.**
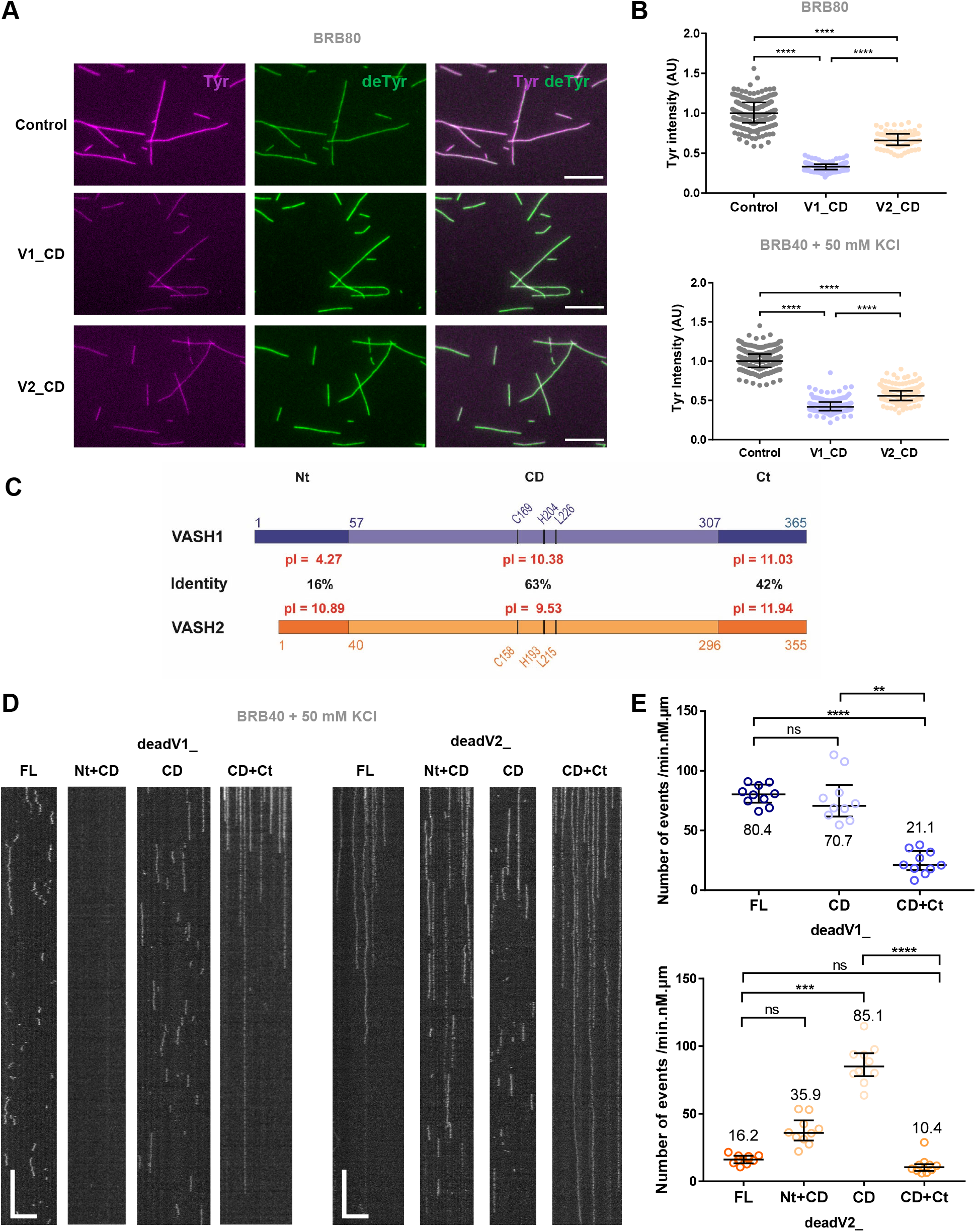
The N- and C-terminal regions of VASH1 and VASH2 are crucial for their microtubule binding and detyrosination activity. **(A, B)** Activity of enzyme complexes containing the sfGFP-tagged central core domains of VASH1 and VASH2 with SVBP (V1_CD and V2_CD) bound to Taxol-stabilized microtubules enriched in HeLa tyrosinated tubulin (Tyr-MTs) measured by immunofluorescence in TIRF experimental conditions (50 pM enzyme, BRB80 or BRB40 supplemented with 50 mM KCl, TIRF chamber). **(A)** Representative images of tyrosinated (Tyr, magenta) and detyrosinated (deTyr, green) α-tubulin pools of microtubules after 30 min in the absence (control) or presence of the enzyme’s core domains. Scale bar, 10 µm **(B)** Analysis of tyrosinated and detyrosinated tubulin signals intensity. Each point represents a microtubule (at least 80 microtubules were analyzed). Data are represented as the median. Statistical significance was determined using Kruskal-Wallis test, ****p < 0,001. **(C)** Schematic representations of human VASH1 (blue) and VASH2 (orange) which are transglutaminase-like cysteine peptidases containing a triad of catalytic residues (Cys, His, and Leu). These proteins share 58% overall sequence identity (74 % homology). Identities between core domains (CD, light colored boxes), and between N- and C-terminal regions (Nt and Ct respectively, dark colored boxes) are specified on the diagram. Isoelectric points (pI) of each domain are also provided. Residues numbers are indicated. **(D, E)** TIRF microscopy study of single molecules of sfGFP-tagged catalytically dead versions of VASH1/2 full-length versions (dead V1_FL and deadV2_FL), core domains (deadV1_CD and deadV2_CD) and truncated versions missing the flanking regions (deadV1_Nt+CD, deadV1_CD+Ct, deadV2_Nt+CD, deadV2_CD+Ct) bound to Taxol-stabilized microtubules enriched in HeLa tyrosinated tubulin (Tyr-MTs). Experiments were performed in the presence of 50 pM enzyme in BRB40 supplemented with 50 mM KCl. **(D)** Representative kymographs. Scale bars: horizontal, 5 µm; vertical, 5 s. **(E)** Analysis of binding frequency. Each point represents a microtubule. Data are represented as the median with the interquartile range. Statistical significance was determined using Kruskal-Wallis test. **p < 0.002 ***p< 0.0002, ****p < 0.0001, and ns, not significant. Results for VASH1-SVBP are in blue and for VASH2-SVBP in orange.

The two VASH core domains were active and, interestingly, they both led to global detyrosination of microtubules (Figure 5AB), as did full-length VASH1-SVBP but not VASH2-SVBP (Figure 1A-C). Analysis of the overall decrease in tyrosinated tubulin signal showed that the VASH2 core domain was significantly less active than the VASH1 core domain under the buffer conditions used. The latter result could be related to the different binding modes of the two complexes with microtubules as observed by cryo-EM: interactions with α- and α‘-tubulin at sites 2 and 3, respectively, could be less favorable for the VASH2 activity than for the VASH1 activity (Figures 3 and 4). As with activity, the complexes containing core domains of VASHs exhibited binding characteristics on microtubules similar to those of the complex with full-length VASH1 (Figure 5DE and Table 3). They displayed residence times of 1.2-1.4 s, thus close to that of full length VASH1-SVBP and at least 10-times shorter than residence time of full length VASH2-SVBP (Table 3). They showed binding frequency close to that of full length VASH1-SVBP and approximately 5-times higher than that of full length VASH2-SVBP (Table 3).

**Table 3.** Binding parameters on microtubules of full-length, truncated, and chimeric versions of VASH1 and VASH2 enzymes in complex with SVBP. **(A)** Related to Figure 5DE. **(B, C)** Related to Figure 6AB. A, B and C represent experiments performed on different days. At least 10 microtubules were analyzed. See Materials and Methods for detailed experimental conditions.

| Experiment (medium) | Ionic strength (mM) | Microtubule | Recombinant protein (concentration) | Residence time $\tau$ (s) | Binding frequency ($\text{min}^{-1} \cdot \text{nM}^{-1} \cdot \mu\text{m}^{-1}$) |
| --- | --- | --- | --- | --- | --- |
| **A** (BRB40, 50 mM KCl) | 140 | Tyr-MTs | deadV1_FL (50 pM) | 1.2 | 80.4 |
|  |  |  | deadV1_Nt+CD (50 pM) | - | No binding |
|  |  |  | deadV1_CD (50 pM) | 1.4 | 70.7 |
|  |  |  | deadV1_CD+Ct (50 pM) | >7.7 | 21.1 |
|  |  |  | deadV2_FL (50 pM) | >17.1 | 16.2 |
|  |  |  | deadV2_Nt+CD (50 pM) | >4.0 | 35.9 |
|  |  |  | deadV2_CD (50 pM) | 1.2 | 85.1 |
|  |  |  | deadV2_CD+Ct (50 pM) | >10.4 | 10.4 |
| **B** (BRB40, 50 mM KCl) | 140 | Tyr-MTs | deadV1_FL (50 pM) | 1.2 | 54.1 |
|  |  |  | deadV1_FL(NtV2) (50 pM) | >9.9 | 31.0 |
|  |  |  | deadV2_FL (50 pM) | >13.4 | 19.1 |
|  |  |  | deadV2_FL(NtV1) (50 pM) | >6.5 | 18.1 |
| **C** (BRB40, 50 mM KCl) | 140 | Tyr-MTs | V1_FL (25 pM) | 1.2 | 96.7 |
|  |  |  | V1_FL(NtV2) (25 pM) | >10.7 | 20.8 |
|  |  |  | V2_FL (25 pM) | >12.9 | 22.9 |
|  |  |  | V2_FL(NtV1) (25 pM) | >3.9 | 42.4 |

Thus, the catalytic core domains of both VASHs behave quite similarly, which is in contrast to their full-length counterparts. Since the strong divergence between the functioning of the two VASH-SVBP enzymatic complexes did not arise from the microtubule-binding mode of these central regions, we hypothesized that their disordered N- and C-terminal flanking regions could explain the observed differences.

### The N- and C-terminal domains of VASHs strongly contribute to the interaction of VASH-SVBP enzyme complexes with microtubules

To understand better the differences in behavior between VASH1-SVBP and VASH2-SVBP, we analyzed the role of their N- and C-terminal disordered regions for interaction of the enzymes with microtubules. To this end, we compared the interaction of dead enzymes containing full-length VASH1/2 (deadV1/2_FL) or their truncated versions lacking either the C-(deadV1/2_Nt+CD) or N-terminal domains (deadV1/2_CD+Ct) with tyrosinated tubulin-enriched microtubules (Tyr-MTs) (Figure 5DE and S1A).

The two mutated enzyme complexes containing the C-terminal region and the catalytic core domain of VASH (CD+Ct proteins) associated strongly to Tyr-MTs, resembling the binding behavior of full-length VASH2-SVBP (Figure 5D). Their mean residence times on microtubules were >7 s (Table 3). In contrast, the two mutant enzymes containing the N-terminal region and the core domain (Nt+CD proteins) behaved very differently: while the VASH1 mutant did not bind to microtubules, the VASH2 mutant showed a long residence-time (Figure 5D and Table 3). Its residence time was even longer than the complex with only the catalytic core domain of VASH2 (τ = 1.2 s for deadV2_CD and τ >4.0 s for deadV2_Nt+CD). Intriguingly, high residence times were usually associated with low binding frequencies, as if long binding events ruled out new ones. This was similarly observed with full-length VASH2-SVBP (Figure 1DF and Table 1A) and could be due to the presence of fluorescence-quenched molecules that were still attached to the microtubule lattice.

Since ionic strength strongly affected both the binding of enzymes to microtubules as well as their detyrosination activity (Figures 2A-D and Table 1), we calculated the isoelectric points of the different regions of VASH1 and VASH2. Interestingly, the C-terminal parts of the VASHs are both very basic, while their N-terminal regions exhibit extremely different isoelectric points. The N-terminal region of VASH1 is acidic as opposed to that of VASH2, which is very basic (Figure 5C). The basic character of the C-terminal parts of VASHs most probably favors binding of the enzyme complexes to the acidic surface of microtubules composed of stretches of acidic glutamates and aspartates from the α- and β-tubulin subunits (Redeker, 2010) (deadV1_CD+CT, deadV2_CD+Ct, Figure 5D and Table 3A). The strong acidic nature of the N-terminal part of VASH1 could explain the lack of binding of the mutant enzyme lacking the basic C-terminal region (deadV1_Nt+CD, Figure 5D and Table 3A). In the case of VASH1-SVBP, its acidic N-terminal part could oppose the interaction of the rest of the protein (core domain and C-terminal region) with microtubules, and thus assist in the release of the enzyme complex from the microtubule. The role of the basic N-terminal part of VASH2 is less clear, but this region might also help the dissociation of the enzyme complex from the microtubule lattice. Indeed, the residence-time of full-length VASH2 is significantly longer than that containing the truncated mutant missing the N-terminal region (τ > 17.1 s for deadV2_FL and τ > 10.4 for deadV2_CD+Ct, Table 3A).

Overall, these results reveal the important contribution of the N- and C-terminal regions of VASH for the interaction of the two enzyme complexes with microtubules. They further suggest an essential role of the VASHs’ N-terminal regions for their divergent activities in vitro, global versus local detyrosination.

### The N-terminal domains of VASHs mediate the divergent microtubule-binding and detyrosination activities

To test the hypothesis that the N-terminal regions of VASHs are critical for the distinct behavior of these enzymes, we constructed chimeras by inverting the N-terminal region of one VASH with the one of the other (Figure S1A). We tested catalytically active and inactive versions of these chimeras for their interaction with microtubules (V1_FL(NtV2), V2_FL(NtV1), deadV1_FL(NtV2), deadV2_FL(NtV1)) and compared their behavior to the ones of the wild type enzymes. Representative kymographs and binding characteristics are presented in Figure 6A-F and Table 3BC. As expected, the presence of the basic N-terminal region of VASH2-SVBP significantly changed the binding behavior of VASH1-SVBP (Videos S1 and S3). The residence-times of the chimeras were more than 8-times higher than those of the native VASH1-SVBP (τ = 1.2 s for V1_FL; τ = 1.2 s for deadV1_FL; τ > 10.7 s for V1_FL(NtV2); τ > 9.9 s for deadV1_FL(NtV2)). Their residence times were therefore more similar to the residence time of full-length VASH2-SVBP (τ > 13.4 s for deadV2_FL; τ > 12.9 for V2_FL). The presence of the acidic N-terminal region of VASH1 also significantly changed the binding behavior of VASH2-SVBP (Videos S2 and S4). The residence time of the chimera on microtubules was more than 4-times shorter than the one of the active wild type enzyme (τ > 12.9 for V2_FL; τ > 3.9 for V2_FL(NtV1)), and 2-times shorter for the dead version (τ > 13.4 for deadV2_FL; τ > 6.5 for deadV2_FL(NtV1)).

**Figure 6.**
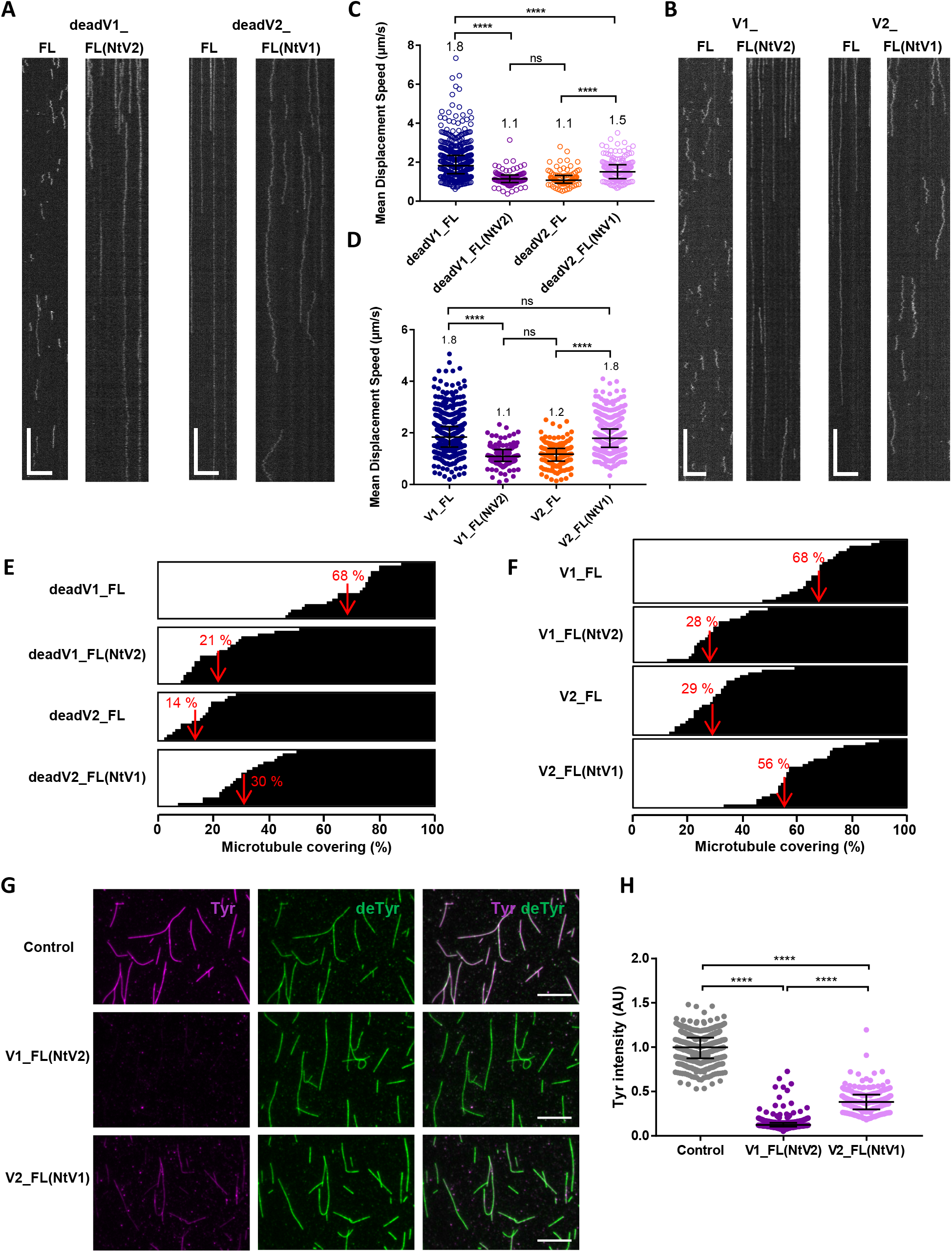
The exchange of the N-terminal regions strongly alters binding behavior and activity of VASH-SVBP enzyme complexes. **(A-F)** Comparison of the interaction with Taxol-stabilized microtubules enriched in HeLa tyrosinated tubulin (Tyr-MTs) of the sfGFP-tagged full length VASH1/2 enzymes (V1_FL, V2_FL) and chimeras in which the N-termini were exchanged (V1_FL(NtV2), V2_FL(NtV1)). Both catalytically dead **(A, C, E)** and active **(B, D, F)** versions were studied. Experiments were performed in the presence of 50 pM enzyme in BRB40 supplemented with 50 mM KCl. **(A, B)** Representative kymographs. Scale bars: horizontal, 5 µm; vertical, 5 s. Examples of TIRF movies from which kymographs were extracted are presented in supplemental data for the chimeras (Videos S3, S4). Binding characteristics are summarized in Table 3. **(C, D)** Analysis of diffusion. Each point represents a single molecule, the molecules moving on at least 17 microtubules were analyzed. **(E, F)** Representation of the microtubule surface covered with VASH-SVBP molecules (white) or not covered (black) during the 45 s of TIRF movies. Each horizontal line represents a microtubule (at least 17 microtubules were analyzed). The red value corresponds to the mean covering (in %). **(G-H)** Activity of VASH-SVBP enzyme complexes and chimeras on Tyr-MTs measured by immunofluorescence in the same experimental conditions (50 pM enzyme, BRB40 supplemented with 50 mM KCl, TIRF chamber). **(G)** Representative images of tyrosinated (Tyr, magenta) and detyrosinated (deTyr, green) α-tubulin pools of microtubules after 30 min incubation in the absence (control) or presence of the indicated enzyme. Scale bar, 10 µM **(H)** Analysis of tyrosinated- and detyrosinated-tubulin signals intensities. Each point represents a microtubule (at least 200 microtubules were analyzed). Data are represented as the median. Statistical significance was determined using Kruskal-Wallis test, ****p < 0.0001.

The exchange of the N-terminal regions also strongly altered the diffusion of the enzyme complexes on microtubules and their covering of the microtubule lattices (Figure 6C-F). The chimeric complex with the N-terminal region of VASH2 and the remainder of VASH1 diffused poorly compared to VASH1-SVBP, and its covering of microtubules was significantly reduced. This was true for both the dead and catalytically active versions of VASH1-SVBP. In contrast, replacement of the N-terminal region of VASH2 with that of VASH1 significantly increased the diffusion and enhanced microtubule covering by VASH2-SVBP. This was particularly evident for the active chimera, which in the presence of the N-terminal region of VASH1 revealed a diffusion capacity equal to that of VASH1-SVBP (Figure 6D). The catalytically dead VASH2 chimera, however, retained a significantly lower diffusion capacity than VASH1-SVBP (and of its active counterpart, Figure 6CD). This difference could be due to the absence of a possible structural change occurring during tyrosine cleavage, which could favor VASH2 release from microtubules. This would be consistent with the significantly lower diffusion capacity of the dead VASH2 version compared to its active counterpart (1.4 µm/s for V2_FL and 1.0 for deadV2_FL in BRB40 supplemented with 50 mM KCl (Figure 1H); 1.6 µm/s for V2_FL and 1.3 for deadV2_FL in BRB80 supplemented with 100 mM KCl, not shown). The binding frequencies of VASH1 chimeras were strongly reduced compared with those of VASH1 (with a 4.7-fold decrease for active versions and a 1.7-fold decrease for dead versions, Table 3BC). The binding frequency was almost unchanged for the dead VASH2 chimera compared to that of the dead VASH2, which was not the case for the active versions, with the VASH2 chimera having a clearly higher binding frequency than VASH2 (1.9-fold increase, Table 3BC).

We then examined the microtubule detyrosination activity of the chimeras (V1_FL(NtV2) and V2_FL(NtV1)) (Figure 6FG) using immunofluorescence. Both chimeras induced global detyrosination, resembling the way VASH1-SVBP acts on microtubules. Thus, the presence of the N-terminal region of VASH1 changed the detyrosinating behavior of VASH2-SVBP from local to global, correlating with the TIRF experiment showing that the chimera is much more diffusive and has a higher binding frequency (Figure 6D, Table 3C). On the other hand, despite inducing an important loss of diffusion capacity (Figure 6D), the N-terminal region of VASH2 did not appreciably switch the behavior of VASH1-SVBP (Figure 6G). The VASH1 chimera still induced global microtubule modifications. This result could be due to the core domain of VASH1 being significantly more active than the one of VASH2 (Figure 5B). Notably, quantification of tyrosinated tubulin levels showed that changes were significantly larger for the VASH1 chimera than for the VASH2 chimera, like for their wild type counterparts.

Together, our data reveal the importance of the N-terminal regions of the VASHs in the divergent binding of VASH-SVBP complexes to the microtubule lattice as well as their detyrosination activity. Indeed, by exchanging these regions, we were able to switch the microtubule-binding behavior and activity of one enzyme to that of the other one.

### VASH1 and VASH2 give rise to distinct detyrosination patterns in cells

To test whether our in vitro results are relevant in cells, we assayed recombinant VASH-SVBP activities on exposed microtubules of murine embryonic fibroblasts (MEFs). Cultured MEFs were incubated in a buffer containing Triton-X100 for lysis and glycerol to stabilize microtubules (see Material and Methods). Figure 7 show that native MEFs mainly contain tyrosinated microtubules and that addition of active VASH-SVBP complexes (V1_FL and V2_FL) at 50 or 200 pM led to a marked increase of detyrosination of cellular microtubules. Most likely due to concentration of microtubules, the observed increase was greater around nuclei. Interestingly and similar to what was observed in vitro, while the VASH1-SVBP complex induced a diffuse and spread microtubule detyrosination, VASH2-SVBP led to the formation of local areas of lattice modifications. Thus, our data show that the two VASH-SVBP enzymes generate different patterns of microtubule detyrosination both in a reconstituted system and in cells.

**Figure 7.**
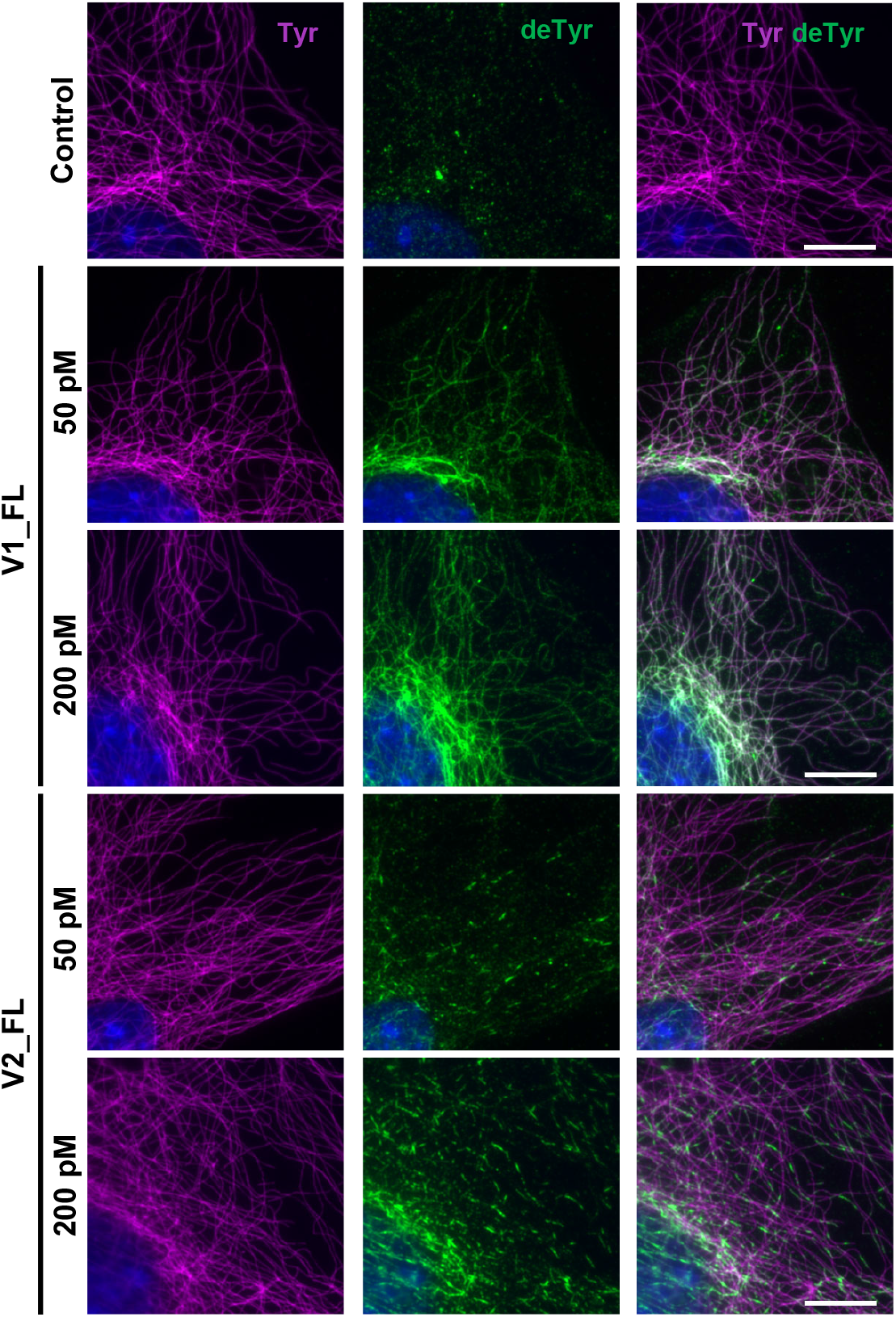
VASH1-SVBP and VASH2-SVBP generate different detyrosination profiles in fibroblast cells. Cultured murine embryonic fibroblasts (MEFs) were permeabilized in BRB80 supplemented with Triton X-100 and glycerol, incubated 30 min with recombinant enzymes (50 or 200 pM, as indicated), fixed in methanol at −20°C and labeled with anti-tyrosinated (Tyr, magenta) and anti-detyrosinated (deTyr, green) tubulin antibodies to reveal detyrosination/tyrosination levels on microtubules. Scale bar, 10 µm.

## DISCUSSION

Our in vitro and in cellulo findings reveal divergent activities of the two VASH-SVBP enzymes, with VASH1-SVBP leading to global microtubule detyrosination and VASH2-SVBP generating local areas of microtubule lattice modifications. These different activities are driven by the different microtubule binding behaviors of the two VASH-SVBP enzymes (see graphical abstract).

We found a moderately stable interaction of the two VASH core domains with the microtubule lattice, which includes the specific recognition of two α-tubulin subunits from adjacent protofilaments of the substrate. While the involved interaction sites confer specificity to this family of tubulin carboxypeptidases, the disordered N- and C-terminal regions of VASHs contribute to their microtubule-binding strength and therefore to their mode of action.

The common C-terminal, strongly positively charged region of the two VASHs considerably lengthen the interaction time of both enzyme complexes with microtubules. A reasonable model explaining this observation is that this basic region interacts through electrostatic attractive interactions with the negatively charged outer surface of microtubules (mostly composed of acidic residues stemming from the C-termini of both α- and β-tubulin (Redeker, 2010)), allowing to hold an enzyme complex close to the microtubule surface as it moves from one tubulin dimer to another. The C-terminal VASH region could thus maintain an enzyme complex bound to the lattice when its catalytic central domain transiently detaches from it upon cleavage of the C-terminal α-tubulin tyrosine to reach the next substrate. Such a mechanism would increase both the residence time of a VASH-SVBP complex with microtubule lattices as well as its activity.

In contrast, the N-terminal domain of the two VASHs could be involved in the release mechanism of the enzyme complex from microtubules, and consequently contribute to its diffusion on the lattice. This is particularly clear for VASH1-SVBP: the VASH1 mutant lacking the N-terminal region shows a significantly increased residence time. Moreover, VASH1-SVBP switches from a diffusive to a static binding behavior on microtubules when the N-terminal region is removed. Due to its many negative charges, this domain probably behaves as an electrostatic “repellent”, promoting detachment of the enzyme from the surface of microtubules. This is not the case for the N-terminal region of VASH2, which in contrast to the one of VASH1 is strongly positively charged; it might thus contribute together with its positively charged C-terminal region to the very long residence time of VASH2-SVBP on microtubules.

Our data suggest that the diffusive capacity of VASH1-SVBP, sustained by its N-terminal domain, promotes the global detyrosination of microtubules by this enzyme in vitro. The local detyrosination observed for VASH2-SVBP likely relies on its weak diffusion capacity along microtubules. The two enzymes can nevertheless be considered as processive, i.e., they are capable to catalyze several consecutive reactions without fully detaching from a microtubule. VASH1-SVBP shows higher detyrosination activity than VASH2-SVBP most probably because it has a higher diffusive capability. In addition, VASH1 might have a higher cleavage efficiency than VASH2, as suggested by the increased detyrosination activity of the central domain of VASH1 compared to the one of VASH2 (Figure 5B).

Notably, our work reveals that as for molecular motors (Zhernov et al., 2020) electrostatic interactions are key factors in the association between VASH-SVBP complexes and microtubules, making their processive enzymatic functioning possible. Indeed, the disordered flanking regions of the highly structured VASH core domains fine-tune the association via such electrostatic interactions. Interestingly, the N-terminal regions of the VASHs bearing opposite charges give rise to both a divergent microtubule-binding behavior as well as a different activity of the two enzyme complexes in a reconstituted system. Other mechanisms, such as specific post-translational modifications (like, e.g., phosphorylation or polyglutamylation) or association of protein partners (such as MAP4 in cardiomyocytes (Yu et al., 2021)) could also positively or negatively modulate these electrostatic interactions in cells, and therefore influence the association between VASH-SVBP and microtubules as well as the enzymes’ detyrosination activity.

In conclusion, our results reveal a novel pathway by which microtubules can be modified differently depending on the detyrosinating enzyme at work. This mechanism could have implications in the diversity of microtubules reported in cells (for a review see (Roll-Mecak, 2019; Roll-Mecak, 2020)), which likely contain different compositions of detyrosinating enzymes. With the very recent discovery of the third detyrosinating enzyme MATCAP (Landskron et al., 2022), which uses a fundamentally different catalytic mechanism (being a metalloprotease instead of a cysteine protease) and adopts an alternative microtubule-binding mode and a different way of detyrosinating microtubules, the complexity of microtubule detyrosination in cells may even be more complex. These different microtubule detyrosinating enzymes likely encode numerous subpopulations of microtubules bearing different tyrosination signal patterns that, when read by effectors, could fine-tune physiological processes in specific regions of the cell. Of particular interest is that the VASH2-SVBP complex, which is expressed weakly in most cells, can generate highly localized spots of detyrosinated tubulin on microtubule. We speculate that these spots, by generating areas on microtubules unable to bind protein partners such as, for example, CAP-Gly proteins or tyrosine-sensitive molecular motors (Chen et al., 2021; Peris et al., 2006; Peris et al., 2009), may have a role in cytoskeletal organization and/or intracellular trafficking in confined regions of cells. Several physiological processes, including cell division, organization, or migration could require such local signals for proper functioning. In this context, a better understanding of the regulation of tubulin-detyrosinating enzymes as well as their spatial and temporal expression patterns in cells becomes essential.

## MATERIAL AND METHODS

### Constructs and enzymes purification for TIRF and immunofluorescence studies

The cDNA encoding human full-length SVBP (NP_955374, residues 1-66) was cloned into the first multiple cloning site of a modified pETDuET-1 vector, to generate a fusion protein with C-terminal myc and FLAG tags. The cDNAs encoding the human VASH1 (NP_055724) and VASH2 (NP_001287985) were cloned into the second multiple cloning site of the modified vector, to generate fusion proteins with an N-terminal superfolder GFP (sfGFP) followed by a PreScission cleavage site, plus C-terminal myc and His tags. The VASH1 and VASH2 constructs used in this study (Figure S1A) were : VASH1 full-length (V1_FL, residues 1-365), dead VASH1 full-length, (deadV1_FL, residues 1-365, C169A), VASH1 core domain (V1_CD, residues 57-307), dead VASH1 core domain (deadV1_CD, residues 57-307, C169A), dead VASH1 core domain + N-ter (deadV1_CD+Nt, residues 1-307, C169A), dead VASH1 core domain + C-ter (deadV1_CD+Ct, residues 57-365, C169A), VASH2 full length (V2_FL, residues 1-355), dead VASH2 full length (deadV2_FL, residues 1-355, C158A), VASH2 core domain (V2_CD, residues 40-296), dead VASH2 core domain (deadV2_CD, residues 40-296, C158A), dead VASH2 core domain + N-ter (deadV2_CD+Nt, residues 1-296, C158A), dead VASH2 core domain + C-ter (deadV2_CD+Ct, residues 40-355, C158A), dead VASH2 with VASH1 N-ter (deadV2_NtV1, VASH1 residues 1-49 and VASH2 residues 40-355, C158A), dead VASH1 with VASH2 N-ter (deadV1_NtV2, VASH2 residues 1-39 and VASH1 residues 51-365, C169A). For the above dead versions, mutations were introduced using a standard PCR procedure. All constructs were verified by DNA sequencing (GENEWIZ). The cloning for non-sfGFP tagged proteins expression were described in (Wang et al., 2019).

Protein expression and purification of the various His-tagged VASH1/2-SVBP complexes were performed as previously described (Wang et al., 2019) with slight modifications. Briefly, BL21(DE3) *E*. *coli* cells were transformed with the corresponding construction and then cultured in Luria-Bertani (LB) medium at 37 °C. Protein expression was induced by addition of 0.5 mM IPTG and overnight incubation at 18 °C. Cells were collected and resuspended in lysis buffer (50 mM HEPES pH 8.0, 500 mM NaCl, 10% glycerol, 20 mM imidazole, 2 mM DTT and cOmplete EDTA-free protease inhibitor cocktail tablets (GE, Healthcare)). Following cell lysis by sonication, the extract was clarified by centrifugation at 100,000 *g* for 30 min at 4 °C. Total lysate was loaded in a HisTrap HP Ni^2+^ Sepharose column (GE Healthcare). Column was then extensively washed using lysis buffer, and enriched VASH1/2-SVBP complexes were eluted from the HisTrap HP Ni^2+^ Sepharose column using elution buffer (lysis buffer with imidazole concentration raised to 400 mM). Protein fractions were further collected and concentrated, and finally purified with a Superdex 200 10/300GL (GE Healthcare) size exclusion column. The concentration of soluble proteins was measured using Bradford assay with bovine serum albumin (BSA) as the standard. All proteins were kept in storage buffer (20 mM Tris-HCl at pH 7.4, 150 mM NaCl, 2 mM DTT) and stored in liquid nitrogen until further use.

### Preparation of tubulin and Taxol-stabilized microtubules

Preparation of brain tubulin and its labelling with either biotin or ATTO-565 fluorophore (ATTO-TEC Gmbh) were performed according to (Ramirez-Rios et al., 2017). HeLa tubulin proteins used in this study were purified as previously described (Souphron et al., 2019). All tubulin proteins were stored in liquid nitrogen.

Taxol-stabilised microtubules were prepared by polymerising 45 µM tubulin (composed of 65% of HeLa tyrosinated or detyrosinated tubulin, 30% biotinylated brain tubulin and 5% ATTO-565-labelled brain tubulin, named Tyr-MT or deTyr-MT, respectively) in BRB80 (80 mM PIPES(piperazine-N,N′-bis(2-ethanesulfonic acid)-KOH at pH 6.8, 1 mM MgCl_2_, 1 mM EGTA) supplemented with 1 mM GTP. Taxol (100 µM) was then added, and microtubules were further incubated for 30 min. Microtubules were then centrifuged for 5 min at 200,000x *g* and resuspended in BRB80/10 µM Taxol buffer. All incubations and centrifugations were carried out at 35°C. Using a method described in (Aillaud et al., 2016), we estimated that Tyr-MTs contained 80 % tyrosinated, 17 % detyrosinated and 3 % Δ2-tubulin, while deTyr-MT contained 82 % detyrosinated, 15 % tyrosinated and 3 % Δ2-tubulin (see Figure S1C).

### In vitro binding assays

Experiments were conducted in the media indicated on Figures: BRB40 or BRB80 (40 or 80 mM PIPES-KOH at pH 6.8, 1 mM MgCl_2_, 1 mM EGTA) buffers supplemented or not with the indicated concentrations of KCl. Ionic force of media was calculated as in (Thiede et al., 2013) (see Table 1). Flow chambers for TIRF imaging assays were prepared as previously described (Stoppin-Mellet et al., 2020). Perfusion chambers were treated with 30 µl neutravidin (Thermofisher, 31000, 25 µg/ml) incubated for 3 min followed by 50 µl PLL20K-G35-PEG2K (0.1 mg/ml, Jenkem) for 30-45 sec. Chambers were then washed 3 times with 100 µl 1% BSA in indicated medium. Subsequently, microtubules enriched in HeLa tyrosinated or detyrosinated tubulin (Tyr-or deTyr-MTs) prepared as above were diluted to 40 µM and perfused into the chamber. Then, the flow chamber was flushed 3 times by 100 µL of 1% BSA/10 µM Taxol in medium to remove unbound microtubules. Finally, sfGFP-VASH1/2-SVBP complexes were perfused in an assay-mix solution (medium supplemented with 82 µg/ml catalase, 580 µg/ml glucose oxidase, 1 mg/ml glucose, 4 mM DTT, 0.5 mM Taxol, 0.017% methylcellulose 1500 cp). All incubations were made at room temperature. The chamber was sealed and then imaged using TIRF microscope.

### TIRF in vitro assay

Images were recorded within the first 30 min following addition of the assay-mix solution (see above) on an inverted microscope (Eclipse Ti, Nikon) equipped with a Perfect Focused System, a CFI Apochromat TIRF 100X/1.49 N.A oil immersion objective (Nikon), a warm stage controller (Linkam Scientific) and a Technicoplast chamber to maintain the temperature, an objective heater (OkoLab), an iLas2 TIRF system (Roper Scientific) and a sCMOS camera (Prime95B, Photometrics) controlled by MetaMorph software (version 7.10.3, Molecular Devices). For dual view imaging, an OptoSplit II bypass system (Cairn Research) was used as image splitter and illumination was provided by 488- and 561-nm lasers (150 mW and 50 mW respectively). Temperature was maintained at 35 °C for all imaging purposes. Acquisition rate was one frame each 50 ms exposure (in streaming acquisition) during 45 s. For each condition at least two slides from two independent experiments using different protein preparations were done.

### In vitro assay of detyrosination activity using immunofluorescence

Immunofluorescence in vitro assays were made using exactly the same perfusion chambers and TIRF experiments conditions as above. After addition of the assay-mix solution, incubation was done at 37°C for 30 min followed by 3 washes with 10 µM Taxol and 1% BSA in indicated medium (wash buffer). Then, incubation with primary antibodies was done for 15 min, followed by 3 washes with 100 µL of wash buffer, incubation with secondary antibodies for 15 min and finally 3 washes with 100 µL of wash buffer. Primary antibodies were rat anti-tyrosinated tubulin (YL1/2 = anti-Tyr, 1:6000) and rabbit anti-detyrosinated tubulin (anti-deTyr, 1:1000) (see (Aillaud et al., 2016)) and secondary antibodies were anti-rat coupled to Alexa Fluor 488 and anti-rabbit coupled to Cyanine 3 (Jackson ImmunoResearch) both diluted to 1:500. Images were obtained using a LEICA DMI600/ROPER microscope controlled by Metamorph Video software using the same illumination conditions. For each condition at least three independent experiments with different protein preparations were done.

### Assays of detyrosination in cells

Murine embryonic fibroblasts (MEFs) were prepa, as previously described (Erck *et al*., 2005). Cells were treated with BRB80 supplemented with 1 % Triton X-100 and 10% glycerol at 35°C for 15 s. Permeabilized cells were washed twice with BRB80 supplemented with 10 % glycerol, incubated with 50 or 200 pM of recombinant sfGFP-VASH-SVBP complexes during 30 min at 37°C, and rinsed twice with BRB80 supplemented with 10 % glycerol.

For immunofluorescence, cells were washed in warm phosphate buffered saline (PBS) supplemented with 0.1 % Tween and fixed in freshly prepared −20°C methanol. They were then incubated with primary antibodies (anti-Tyr and anti-deTyr; 1:500), followed by incubation with secondary antibodies (anti-rat conjugated with Alexa Fluor 488 ; anti-rabbit conjugated with Cyanine 3 ; 1:500). Samples were mounted using DAKO medium supplemented with Hoescht 33258 (1 µg/ml). Images were captured with a Zeiss Axiovert 200M microscope equipped with the acquisition software MetaMorph using the same illumination conditions.

### SDS-PAGE, immunoblotting and antibodies

Equal amounts of purified proteins (1.5 µg) were loaded onto 4-20% gradient polyacrylamide gels (Biorad) and separated by electrophoresis. Proteins were transferred to nitrocellulose membranes and probed as follow: rabbit anti-GFP from Life Technologies (1:10,000), mouse anti-His from Qiagen (1:10,000), home-made rabbit anti-VASH1, anti-VASH2 and anti-SVBP (all used at 1:10,000), and mouse anti-α tubulin (αtot, 1:5,000), rat anti-Tyr (YL_1/2_, 1:5,000), rabbit anti-deTyr (1:20,000), rabbit anti-Δ2 tubulin (1:20,000). All anti-tubulin antibodies were described in (Aillaud et al., 2016). Anti-VASH1 Gre was described in (Chen et al., 2020). Anti-VASH2 Gre and anti-SVBP Gre were produced in a similar manner against peptides AIRNAAFLAKPSIPQVPNYRLSMTI of VASH2 and DPPARKEKSKVKEPAFRVEKAKQKS of SVBP, respectively. Incubation with primary antibodies were followed by incubation with secondary antibodies conjugated with horseradish peroxidase (Jackson ImmunoResearch). Blots were finally revealed using Pierce™ ECL Western Blotting Substrate (ThermoFisher Scientific) and a ChemiDoc MP Imaging System (BioRad) according to the manufacturer’s protocol.

### Cloning and protein purification of VASH2-SVBP sample for cryo-EM studies

The human cDNAs of full length VASH2 (UniProtKB ID Q86V25) and full length SVBP (Q8N300) were codon optimized for *E. coli* expression and synthesized (GENEWIZ). The sequence of VASH2 was modified by mutating Cys158 to alanine resulting in a catalytically inactive protein (Wang et al., 2019). The synthetic cDNAs were then cloned into two different plasmids, VASH2 into pET3d (Novagen) and SVBP into pET28 (EMD Biosciences). When inserted into the pET28 vector, the SVBP sequence was fused to a His_6_ tag at its C-terminus for purification purposes. Protein expression and purification of the His-tagged VASH2-SVBP complex was performed as previously described (Wang et al., 2019). Final protein samples were snap frozen in liquid nitrogen at 2 mg/ml concentration and stored at −80°C.

GMPCPP-stabilized microtubules were prepared using HeLa cell tubulin prepared as in (Souphron et al., 2019). Purified tubulin was resuspended at 4 mg/ml on ice using BRB80 buffer and GMPCPP was added to 0.5 mM final concentration. The tubulin solution was incubated on ice for 5 min then centrifuged for 10 min at 16,000x g at 4°C. The supernatant was transferred to a fresh tube and placed on a 37°C heat block for 40 min. Subsequently, the microtubule solution was kept at 20°C before use.

### Cryo-EM grid preparation

Glow discharged Quantifoil Cu300 R2/1 holey carbon EM grids (Electron Microscopy Sciences) were used for specimen preparation. Microtubules were centrifuged for 10 min at 16,000x *g* and thoroughly resuspended in BRB80 buffer to 2 mg/ml. 3.5 μl of the microtubule solution was applied to the EM grid and incubated for 1 min at room temperature. The grid was blotted manually and 3.5 μl of a 2 mg/ml V2-SVBP solution was applied to the grid that was then transferred immediately to the chamber of the Vitrobot (ThermoFisher) pre-equilibrated at 25°C and 100% relative humidity. After 1 min of incubation, the grid was blotted for 1 s at blot force of 0 and plunge frozen into liquid ethane. Grids prepared this way were stored in liquid nitrogen prior to imaging.

### Cryo-EM data collection and image processing

Movies were recorded on a Titan Krios at 300 keV (Thermo Fisher), with a GIF Quantum LS Imaging filter (20 eV slit width) and a K2 Summit electron counting direct detection camera (Gatan) at a magnification of 47,260 x, resulting in a pixel size of 1.058 Å, using SerialEM (Mastronarde, 2005). The defocus varied between –0.6 and –2.2 µm. For the Hela sample, 3,345 movies were recorded with a total dose of 65 e-/Å^2^ per movie (9 sec exposure in total, 0.2 sec per frame, 45 frames in total). The dose rate was ∼7.2 e-/Å^2^ per second (∼1.4 e-/Å^2^ per frame). For image processing, the software Relion 3.0.8 (Scheres, 2012) was used. The protocols described in (Cook et al., 2020) and (Debs et al., 2020) were followed with some adaptations. In brief, the movies were drift-corrected and dose-weighted using MotionCor2 (Zheng et al., 2017) and the contrast transfer function parameters were estimated from the motion-corrected electron micrographs using Gctf (Zhang, 2016). Afterwards, microtubule sections were manually picked from the HeLa tubulin data set. Boxes with a size of 600 pixels, re-scaled to 150 pixels with inter-box distance of 82 Å, were extracted. Superparticles were created and processed as previously described (Cook et al., 2020). To compensate for the incomplete decoration of VASH2-SVBP to the microtubule in our cryo-EM sample and improve the signal-to-noise of the VASH2-SVBP density, we selected and subtracted regions of the protofilament where VASH2-SVBP had bound to generate the final map. After subtraction of all possible VASH2-SVBP binding sites at α-tubulin, 3D classification of the particles was performed to identify sites where VASH2-SVBP had bound to the microtubule in a manner similar to described previously (Debs et al., 2020). Single-particle refinement of the particles containing VASH2-SVBP decorated tubulin was performed to generate the final structure.

### Model building and refinement

An initial structural model of the microtubule bound VASH2-SVPB complex was generated in Chimera by combining the GMPCPP-stabilized microtubule (Zhang, 2016) (PDB ID 3JAL) with the crystal structure of the human VASH2-SVBP complex (PDB ID 6QBY). The models were imported into Chimera (Pettersen et al., 2004), and the Fit-in-Map function was used to perform a rigid-body fit of one VASH2-SVBP complex structure and two laterally aligned αβ-tubulin heterodimers into the final cryo-EM map. The models were then combined and refined in Coot (Emsley and Cowtan, 2004) using ‘Set Geman-McClure alpha 0.01’ while all molecule self-restraints were set to 5.0 in the ProSMART module (Nicholls et al., 2014). The model was then further refined in Coot using the ‘Chain Refine’ command. Finally, the model and map were exported to Phenix (Liebschner et al., 2019) and the ‘Real-space Refinement’ tool was used to complete the refinement. MolProbity (Chen et al., 2010) was used to assess the model quality and generate the values reported in Table 2. Chimera and PyMOL (Schrodinger, 2020) were used for the generation of figures.

### Statistical analysis

#### Data analysis of TIRF

Binding/tracking of single VASH1/2-SVBP molecules for estimation of binding parameters and diffusion were measured on kymographs using FIJI software and a homemade plugin KymoTool (Ramirez-Rios et al., 2016). The tracks were detected from kymographs using “Detection plugin”. Briefly, kymograph images were enhanced by substracting a blurred image and convolved for enhancement of continuous tracks. After thresholding, the detected tracks were skeletonized and the tracks composed of several molecules were cleaned manually.

Mean residence time was calculated by fitting the data points to a plateau followed by one-phase decay using GraphPad Prism 7. A precise determination of the residence time for some proteins (such as full-length VASH2-SVBP) was not possible in some experimental conditions and was therefore expressed as “superior to”. Indeed, the molecules were already attached to microtubules at the beginning of the TIRF movies and single molecule traces disappeared during movie acquisition. This latter observation was due to quenching of the sfGFP fluorophore, since at lower intensity of the excitation lamp we observed much longer binding events (not shown). However, at these lower lamp intensities, the signal-to-noise ratio was too low for correct analysis of binding characteristics.

Binding frequency calculations were determined by dividing the number of bindings events by the duration of the recording (in min) and by nanomolar VASH1/2-SVBP protein concentration over the microtubule size (in µm).

For diffusion analysis, each track was interpolated with 1 pixel interval and (X, Y) coordinates were collected. The total displacement (sum of dX at each Y) was calculated, and the final displacement (Xfinal - X0) and time of binding (Yfinal - Y0) were calculated. The displacement speed was then estimated as (total displacement in µm)/(binding time in seconds); thus, it included the displacement of the molecule in both microtubule directions.

For quantification of the surface covered by sfGFP (VASH-SVBP complexes) on microtubule, a binary image was produced for each kymograph. The kymograph was resliced to get a stack where different slices represent the time dimension. A maximal projection was generated and the percentage of the image covered by sfGFP was measured. After independently analyzing each kymograph, a scheme was created where each line represents the percentage of sfGFP signal covering on one microtubule.

The significance of difference among multiple groups was tested by non-parametric ANOVA with the Kruskal-Wallis test, and post-hoc pairwise comparisons were tested by the Dunn’s test except for Figure 1C where the Conover-Iman test was employed. For the interpretation of the p values, ‘ns’ means there is no significant difference between the two distributions. * means p value < 0.05, ** means p value < 0.01, *** means p value < 0.001, and **** mean p value < 0.0001. Data are generally represented as the median with the interquartile range.

#### Data analysis of IF

For measurement of tyrosinated- and detyrosinated tubulin (Tyr, deTyr) intensities FIJI software was used. A mask was created by addition of the two channels, and each object in the image was saved as a region of interest (ROI). The mean intensity of ROIs was measured on each channel after background subtraction. In each experiment, the values were normalized to the median of the control without enzymes. Data analysis was then performed with GraphPad Prism 7. Statistical analyses were obtained using nonparametric one-way ANOVA assuming non-paired and non-Gaussian distribution (Kruskal-Wallis test). For the interpretation of the p values, ns means there is no significant difference between the two distributions, ** means p value < 0.01, and **** mean p value < 0.0001. Data are generally represented as the median with the interquartile range.

## ONLINE SUPPLEMENTAL MATERIAL

**Figure S1.** Proteins used in the TIRF experiments.

**Figure S2.** Tubulin detyrosinating activity of sfGFP-tagged and untagged VASH-SVBP complexes and residence time analysis over the course of a TIRF experiment.

**Figure S3.** Cryo-EM data collection and processing.

**Figure S4**. Sequence alignments of VASHs.

**Figure S5.** Relative position of the catalytic site in microtubule-VASH1 and -VASH2 complexes with respect to the C-terminus of helix H12 of a-tubulin.

**Video S1**. Interaction of single molecules of sfGFP-VASH1-SVBP (V1_FL) with Taxol-stabilized microtubules enriched in tyrosinated HeLa tubulin. Scale bar, 2 μm. **VASH1 complex exhibited short and frequent binding events, and diffused in both directions.**

**Video S2.** Interaction of single molecules of sfGFP-VASH2-SVBP (V2_FL) with Taxol-stabilized microtubules enriched in tyrosinated HeLa tubulin. Scale bar, 2 μm. **VASH2 complex bound less frequently, for much longer times, and diffused significantly less on microtubules than VASH1 complex**.

**Video S3.** Interaction of single molecules of chimeric sfGFP-VASH1-SVBP complex bearing the N-terminal region of VASH2 (V1_FL(NtV2)) with Taxol-stabilized microtubules enriched in tyrosinated HeLa tubulin. Scale bar, 2 μm. The presence of the basic N-terminal region of VASH2-SVBP significantly changed the binding behavior of VASH1-SVBP. The chimeric complex showed much higher residence time and diffused poorly along the microtubule, resembling full length VASH2 complex.

**Video S4**. Interaction of single molecules of chimeric sfGFP-VASH2-SVBP complex bearing the N-terminal region of VASH1 (V2_FL(NtV1)) with Taxol-stabilized microtubules enriched in tyrosinated HeLa tubulin. Scale bar, 2 μm. The presence of the acidic N-terminal region of VASH1 strongly changed the behavior of VASH2-SVBP, with shorter residence time and recovery of diffusion capacity, resembling full-length VASH1 complex.

## Supporting information

Graphical abstract

## ACKNOWLEDGMENTS

We would like to thank Isabelle Jacquier and Lisa De Macedo of Grenoble Institute Neuroscience (GIN) for technical help in preparation of proteins, Yasmina Saoudi of PIC-GIN for helping us with TIRF microscopy, and Benoît Decherf for the design of the FIJI software plugin for detection of single molecule tracks in the kymographs of TIRF experiments. We thank Yves Goldberg for the valuable discussion on statistics. Modified plasmid pETDUet-1 was a generous gift from Sandra Jeudy (IGS, Marseille). We thank the Electron Microscopy Facility (EMF) of the Paul Scherrer Institute and the BioEM Lab of the Biozentrum, University of Basel, for excellent access to cryo-electron microscopes.

This work was supported by the Leducq Foundation, Research grant n° 20CVD01 (to MJM), by Agence National de la Recherche (ANR), grant SPEED-Y n° ANR-20-CE16-0021 (to MJM), and ANR-20-CE13-0011, (to CJ), by the France Alzheimer grant 2023 (to MMM), by Institut National de la Santé et de la Recherche Médicale (INSERM), Centre National de la Recherche Scientifique (CNRS), and University Grenoble Alpes, and the Swiss National Science Foundation (SNSF), grant n° 310030_192566 (to MOS). Part of the salaries of CS and SRR were from a collaborative program between Servier laboratories and MJM team. This work was supported by the Photonic Imaging Center of Grenoble Institute Neuroscience (PIC-GIN, Univ Grenoble Alpes – Inserm U1216) which is part of the ISdV core facility and certified by the IBiSA label.

## Data deposition

The cryo-EM electron density map and the structural model of the microtubule-VASH2-SVBP complex have been deposited in the Electron Microscopy Data Bank (EMD-14634) and the Protein Data Bank (PDB ID 7ZCW), respectively.

## AUTHOR CONTRIBUTIONS

CB analyzed sequences. BB and CB designed and performed cloning experiments for the production of sfGFP-tagged proteins. SRR, CS, and SRC prepared recombinant proteins. SRR and CS conceived and performed single molecule TIRF experiments and assays of activity by immunofluorescence in reconstituted system and in cells. CS generally analyzed binding characteristics (TIRF), and SRR enzyme activity (immunofluorescence) and diffusion (TIRF). ED developed ImageJ macros for TIRF (diffusion) and immunofluorescence analysis, and helped in statistical analysis. FC started the TIRF project with the help of JMS. VSM and IA provided essential advices for single molecule TIRF all along the project. SRR and CS produced, purified and labelled bovine brain tubulin with the help of IA and VSM. MMM and CJ generated and purified HeLa tyrosinated and detyrosinated tubulin. SRC prepared cryo-EM grids. TB collected cryo-EM data. TB and SRC processed and refined cryo-EM data. SRC modelled cryo-EM data. SRC and MOS performed structural analyses. MJM supervised the functional experiments. MOS supervised the structural experiments. MJM prepared Figures 1, 2, 5, 6, S1, S2, Tables 1, 3 and graphical abstract with the help of SRR and CS. SRC prepared Figures 3, 4, S3, S4, S5 and Table 2. SRR prepared Figure 7 and supplemental movies. MJM wrote the manuscript with contributions from all authors.

