## Supplementary figures and images for "VASH1-SVBP and VASH2-SVBP generate different detyrosination profiles on microtubules"

### Graphical abstract

#### GRAPHICAL ABSTRACT

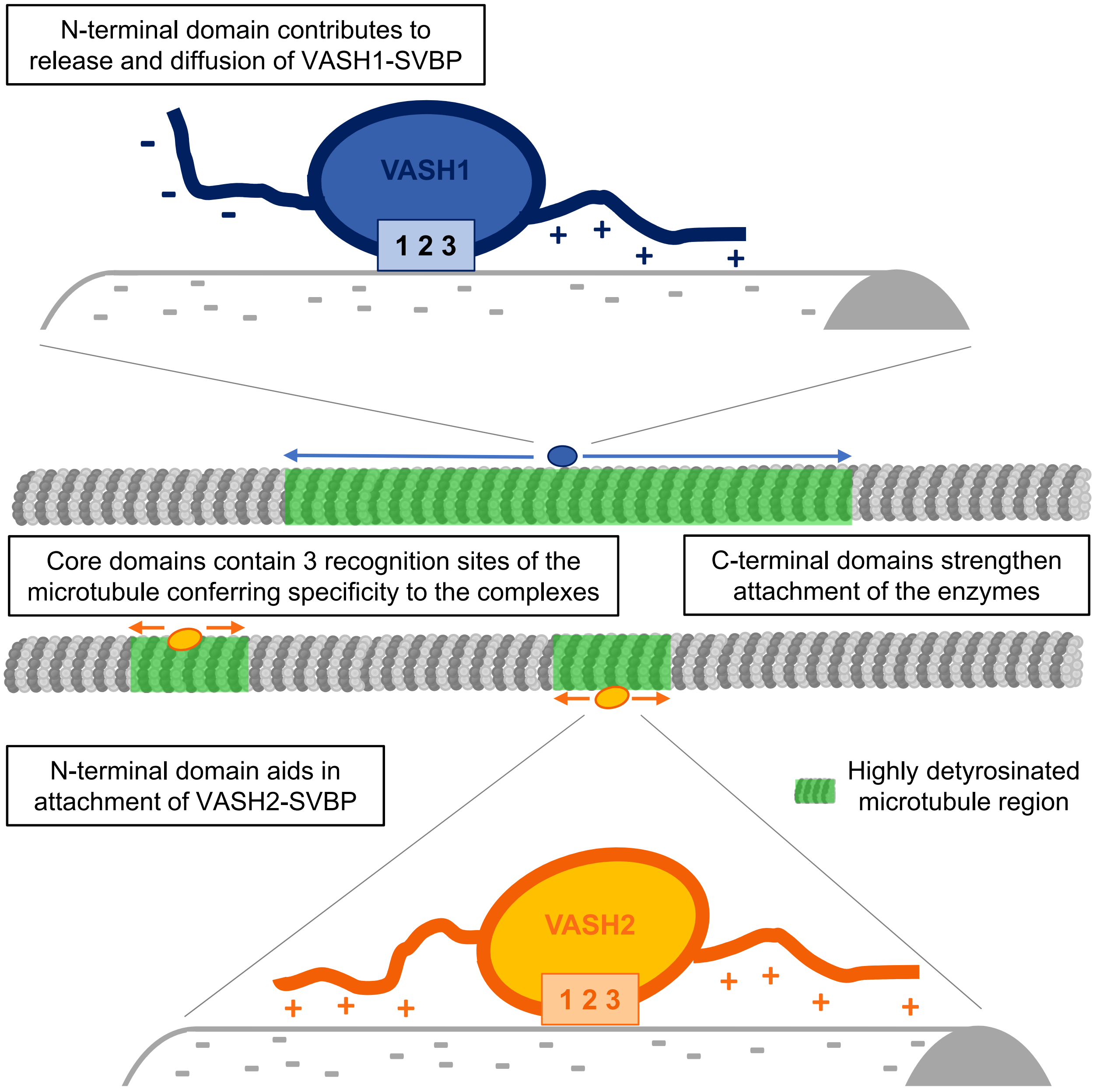
