## supplemental data for "VASH1-SVBP and VASH2-SVBP generate different detyrosination profiles on microtubules"

### SUPPLEMENTAL MATERIALS

Figure S1. **Proteins used in the TIRF experiments**. **(A)** Schematic view of the proteins with color code as in Figure 5. VASH1 (V1) is represented in blue, VASH2 (V2) in orange, SVBP in purple, sfGFP (superfolder GFP) in green, tags (myc, FLAG, His) in gray. Vertical bars, catalytic triads. FL, full length; CD, core domain; Nt, N-terminal region; Ct, C-terminal region. For most proteins, an inactive protein (dead version, with the catalytic cysteine residue mutated into an alanine residue) was also used. **(B)** SDS-PAGE and immunoblots of a complete set of VASH1-SVBP protein preparations. **(C)** Immunoblot of three separate brains tubulin preparations (1, 2, 3), and of tyrosinated- and detyrosinated-HeLa tubulin preparations (Tyr, deTyr). These samples were co-analyzed with extracts from HEK293T cells transfected with various mCherry α-tubulin variants (Tyr, deTyr, Δ2). The mean content of the different forms of α-tubulin present in a brain tubulin preparation was estimated after normalization to total α-tubulin levels (and antibody sensitivity (as in (Aillaud et al., 2016)) to be of 43 %, 49.5 % and 7.5 %, for tyrosinated-, detyrosinated- and Δ2-tubulin respectively. HeLa tubulin was either 100 % tyrosinated or 100 % detyrosinated. Taxol stabilized microtubules enriched either in tyrosinated- or detyrosinated-HeLa tubulin (Tyr-MTs and deTyr-MTs) were composed of 65% of HeLa tubulin and 35 % of brain tubulin (see Materials and Methods).

Figure S2. **Tubulin detyrosinating activity of sfGFP-tagged and untagged VASH-SVBP complexes (A-E) and residence time analysis over the course of a TIRF experiment (F).** VASH1/2-SVBP proteins (V1_FL and V2_FL) were produced with or without an sfGFP tag (see Materials and Methods) and activity of 50 pM of the different complexes was measured by immunofluorescence in BRB40 with 50 mM KCl on Tyr-MTs. **(A, C)** Representative images of tyrosinated and detyrosinated α-tubulin pools of microtubules after 30 min incubation in the absence (control) or presence of the indicated enzyme complexes. Scale bar, 10 µm. **(B, D)** Analysis of tyrosinated-tubulin signals intensities of the experiments presented in A and C, respectively. Each point represents a microtubule (at least 150 microtubules were analyzed). Data are represented as the median with the interquartile range. Kruskal-Wallis test, ****p < 0.0001 and ns, not significant. **(E)** Analysis over time of the decrease in intensity of the tyrosinated-tubulin signals. Each point represents a microtubule (at least 115 microtubules were analyzed). Data are represented as the median with the interquartile range. Statistical significance was determined using Kruskal-Wallis test, ****p < 0.0001. **(F)** Residence time analysis of active and dead versions of VASH1-SVBP during a TIRF experiment performed the same day and in the same experimental conditions as in E. Each point represents the mean residence time of enzymes moving on at least 4 microtubules (1 or 2 slides were analyzed per indicated time).

Figure S3. **Cryo-EM data collection and processing. (A)** Representative cryo-EM micrograph of microtubule-VASH2-SVBP specimens. (**B)** Fourier Space Correlation (FSC) resolution plots of the microtubule-VASH2-SVBP reconstruction. (**C)** Local resolution map of the microtubule-VASH2-SVBP reconstruction. (**D, E**) Exerts of the microtubule-VASH2-SVBP electron density map highlighting the quality of representative regions of αβ-tubulin, VASH2, and nucleotides.

Figure S4. **Sequence alignments of VASHs. (A)** ClustalX multiple sequence alignment of VASH2: *H. sapiens* (NP_001287985); *M. musculus* (NP_659128); *G. gallus* (XP_015139368); *D. rerio* (XP_005160797); *X. laevis* (XP_018118075). Black triangles highlight microtubule-binding residues of VASH2 (Figure 3). **(B)** ClustalX sequence alignment of human VASH1 and human VASH2. Black triangles highlight residues implicated in microtubule interaction of VASH1 (Li et al., 2020) and VASH2 (Figure 3).

Figure S5. **Relative position of the catalytic site in microtubule-VASH1 and -VASH2 complexes with respect to the C-terminus of helix H12 of a-tubulin.** The centroid of the catalytic residue triad is displaced by 6.3 Å in VASH1 (Li et al., 2020) compared to VASH2 due to the observed tilt between the two enzyme complexes (Figure 4). However, the distance between the catalytic triad centroid of VASH1 and VASH2 to the Cα atom of residue S439 of helix H12 of a-tubulin is very similar (29.7 versus 27.3 Å, respectively).

**Video S1**. Interaction of single molecules of sfGFP-VASH1-SVBP (V1_FL) with Taxol-stabilized microtubules enriched in tyrosinated HeLa tubulin. Scale bar, 2 µm. **VASH1 complex exhibited short and frequent binding events, and diffused in both directions.**

**Video S2.** Interaction of single molecules of sfGFP-VASH2-SVBP (V2_FL) with Taxol-stabilized microtubules enriched in tyrosinated HeLa tubulin. Scale bar, 2 µm. **VASH2 complex bound less frequently, for much longer times, and diffused significantly less on microtubules than VASH1 complex**.

**Video S3.** Interaction of single molecules of chimeric sfGFP-VASH1-SVBP complex bearing the N-terminal region of VASH2 (V1_FL(NtV2)) with Taxol-stabilized microtubules enriched in tyrosinated HeLa tubulin. Scale bar, 2 µm. The presence of the basic N-terminal region of VASH2-SVBP significantly changed the binding behavior of VASH1-SVBP. The chimeric complex showed much higher residence time and diffused poorly along the microtubule, resembling full length VASH2 complex.

**Video S4**. Interaction of single molecules of chimeric sfGFP-VASH2-SVBP complex bearing the N-terminal region of VASH1 (V2_FL(NtV1)) with Taxol-stabilized microtubules enriched in tyrosinated HeLa tubulin. Scale bar, 2 µm. The presence of the acidic N-terminal region of VASH1 strongly changed the behavior of VASH2-SVBP, with shorter residence time and recovery of diffusion capacity, resembling full-length VASH1 complex.

Figure S1

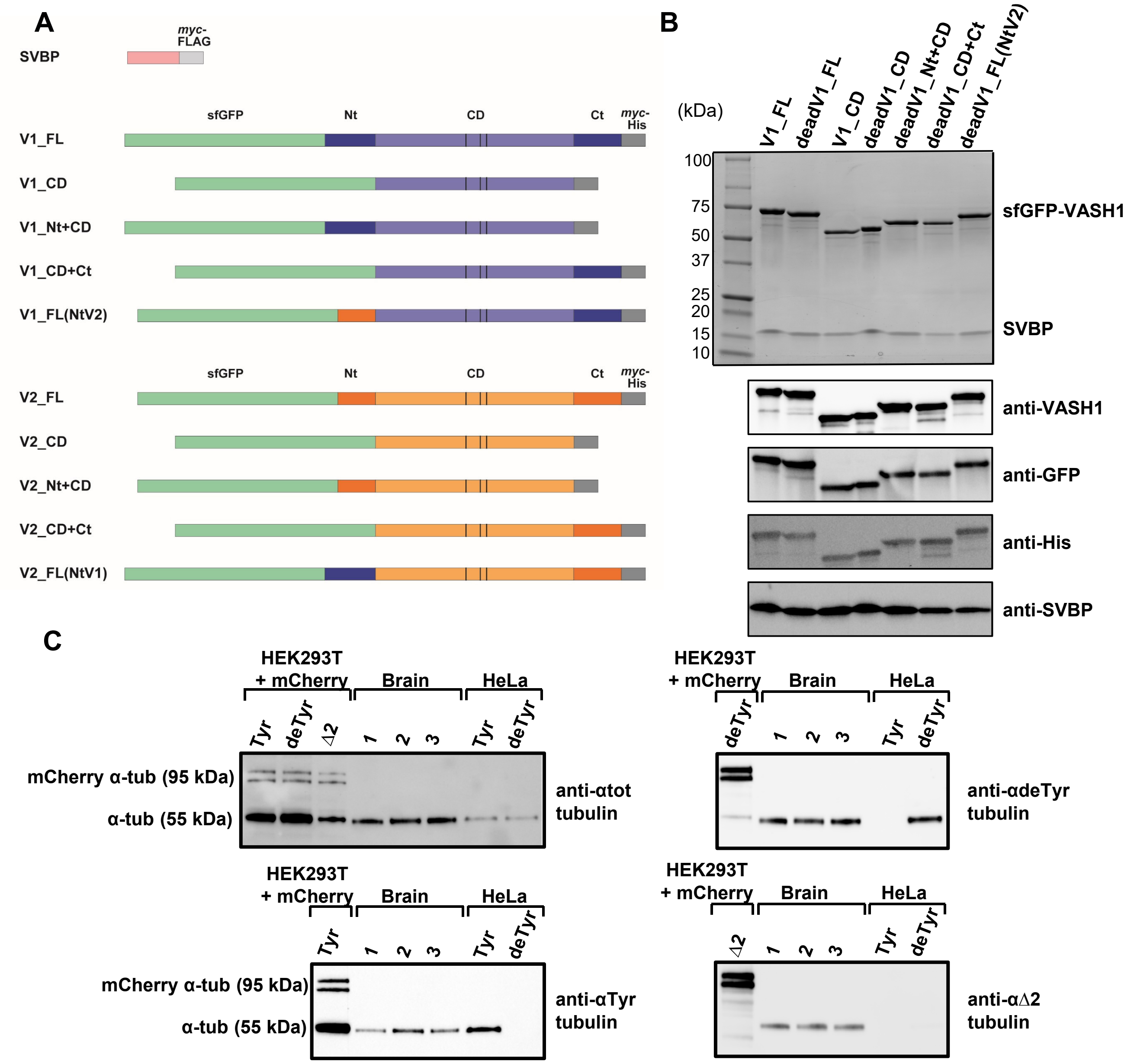

A
SVBP
myc-FLAG
sfGFP
Nt
CD
Ct
myc-His
V1_FL
V1_CD
V1_Nt+CD
V1_CD+Ct
V1_FL(NtV2)
V2_FL
V2_CD
V2_Nt+CD
V2_CD+Ct
V2_FL(NtV1)
B
(kDa)
V1_FL
deadV1_FL
V1_CD
deadV1_CD
deadV1_Nt+CD
deadV1_CD+Ct
deadV1_FL(NtV2)
100
75
50
37
25
20
15
10
sfGFP-VASH1
SVBP
anti-VASH1
anti-GFP
anti-His
anti-SVBP
C
HEK293T + mCherry
Brain
HeLa
Tyr
deTyr
Δ2
1
2
3
mCherry α-tub (95 kDa)
α-tub (55 kDa)
anti-αtot tubulin
anti-αdeTyr tubulin
anti-αTyr tubulin
anti-αΔ2 tubulin

### Figure S2

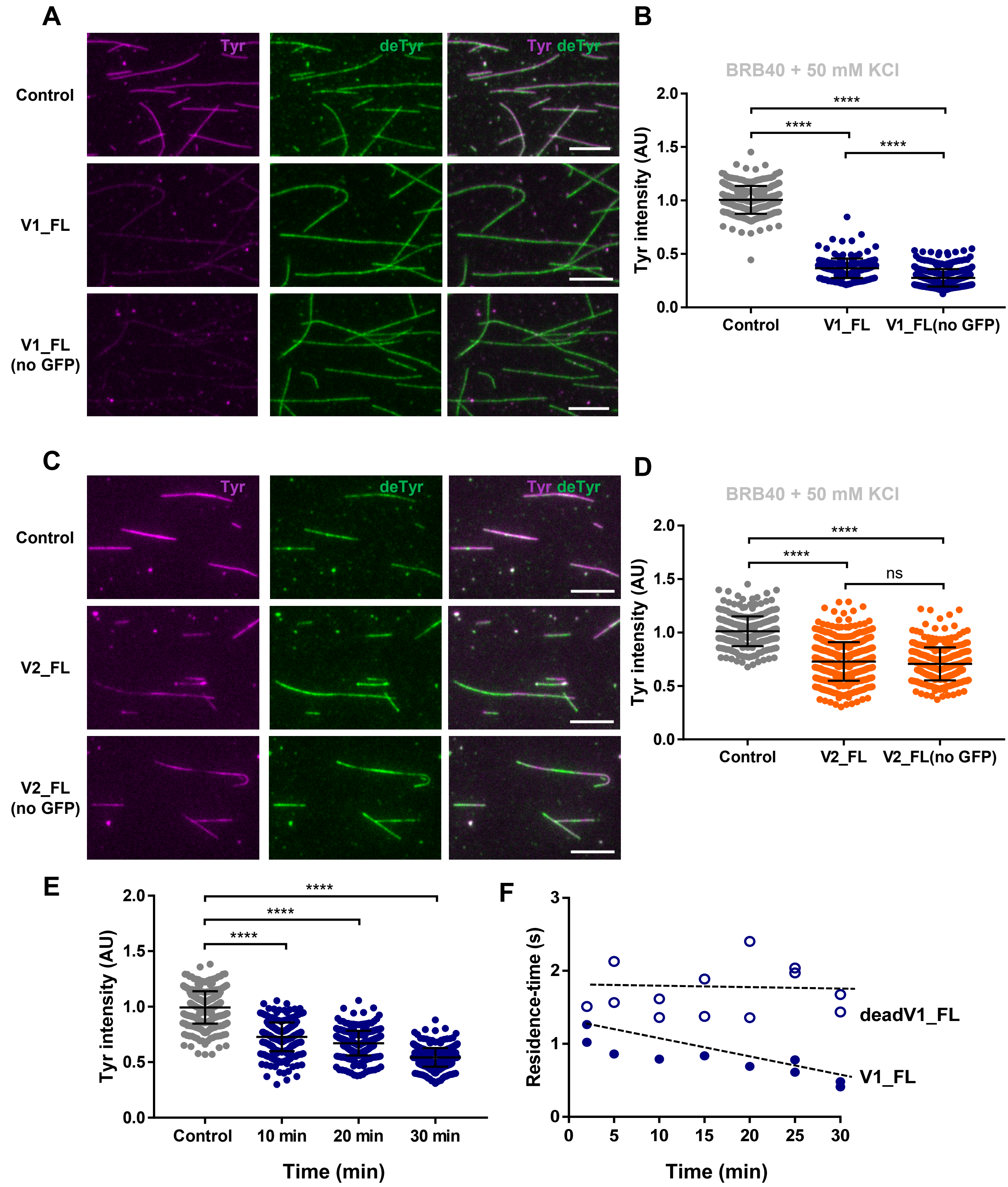
A
Tyr
deTyr
Tyr deTyr
Control
V1_FL
V1_FL
(no GFP)
B
BRB40 + 50 mM KCl
Tyr intensity (AU)
2.0
1.5
1.0
0.5
0.0
****
****
****
Control
V1_FL
V1_FL(no GFP)
C
Tyr
deTyr
Tyr deTyr
Control
V2_FL
V2_FL
(no GFP)
D
BRB40 + 50 mM KCl
Tyr intensity (AU)
****
****
ns
Control
V2_FL
V2_FL(no GFP)
E
Tyr intensity (AU)
****
****
****
Control
10 min
20 min
30 min
Time (min)
F
Residence-time (s)
3
2
1
0
deadV1_FL
V1_FL
0
5
10
15
20
25
30
Time (min)

Figure S3

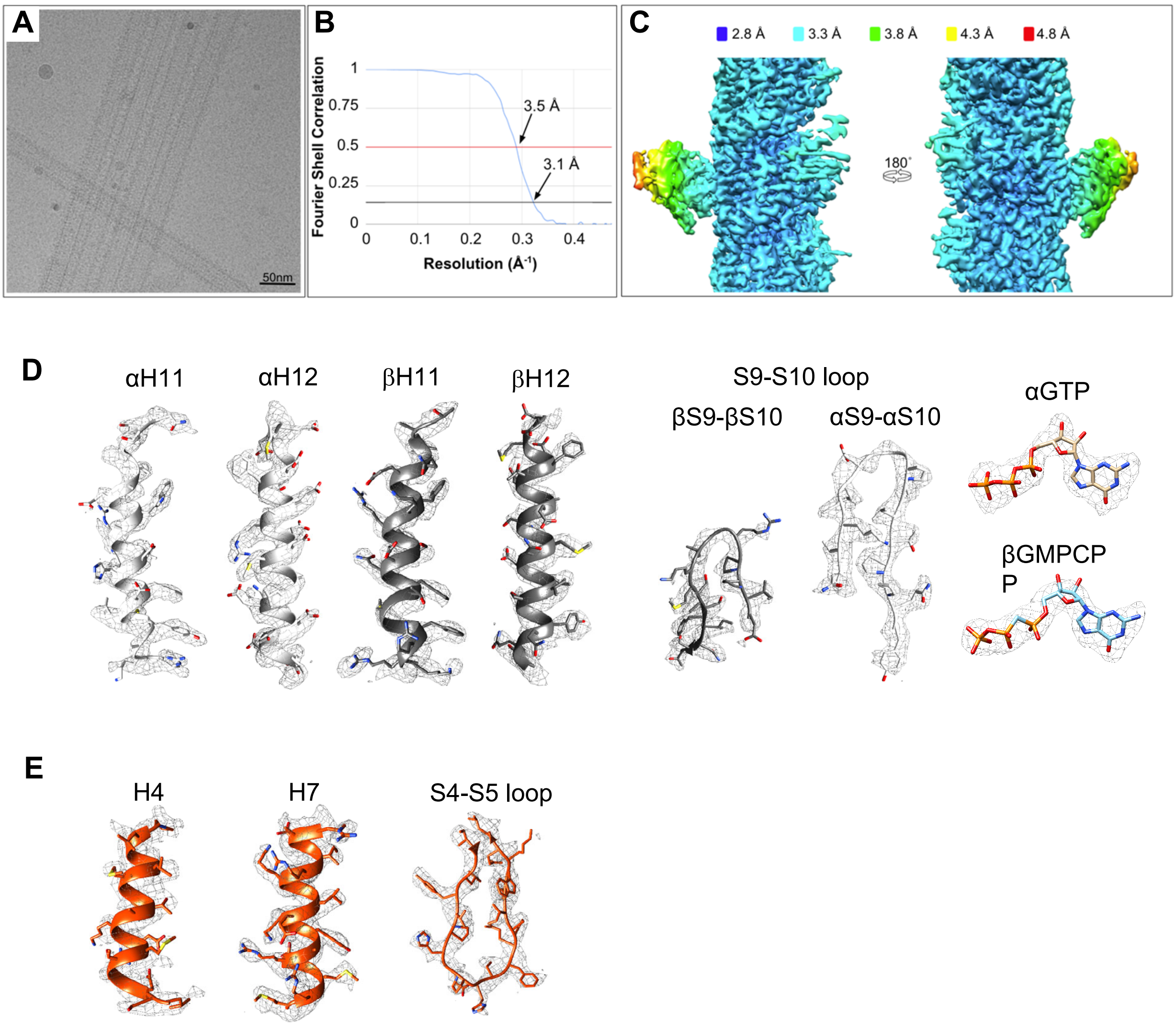

A
50nm
B
Fourier Shell Correlation
1
0.75
0.5
0.25
0
0.1
0.2
0.3
0.4
3.5 Å
3.1 Å
Resolution (Å⁻¹)
C
2.8 Å
3.3 Å
3.8 Å
4.3 Å
4.8 Å
180°
D
αH11
αH12
βH11
βH12
S9-S10 loop
βS9-βS10
αS9-αS10
αGTP
βGMPCP
P
E
H4
H7
S4-S5 loop

Figure S4

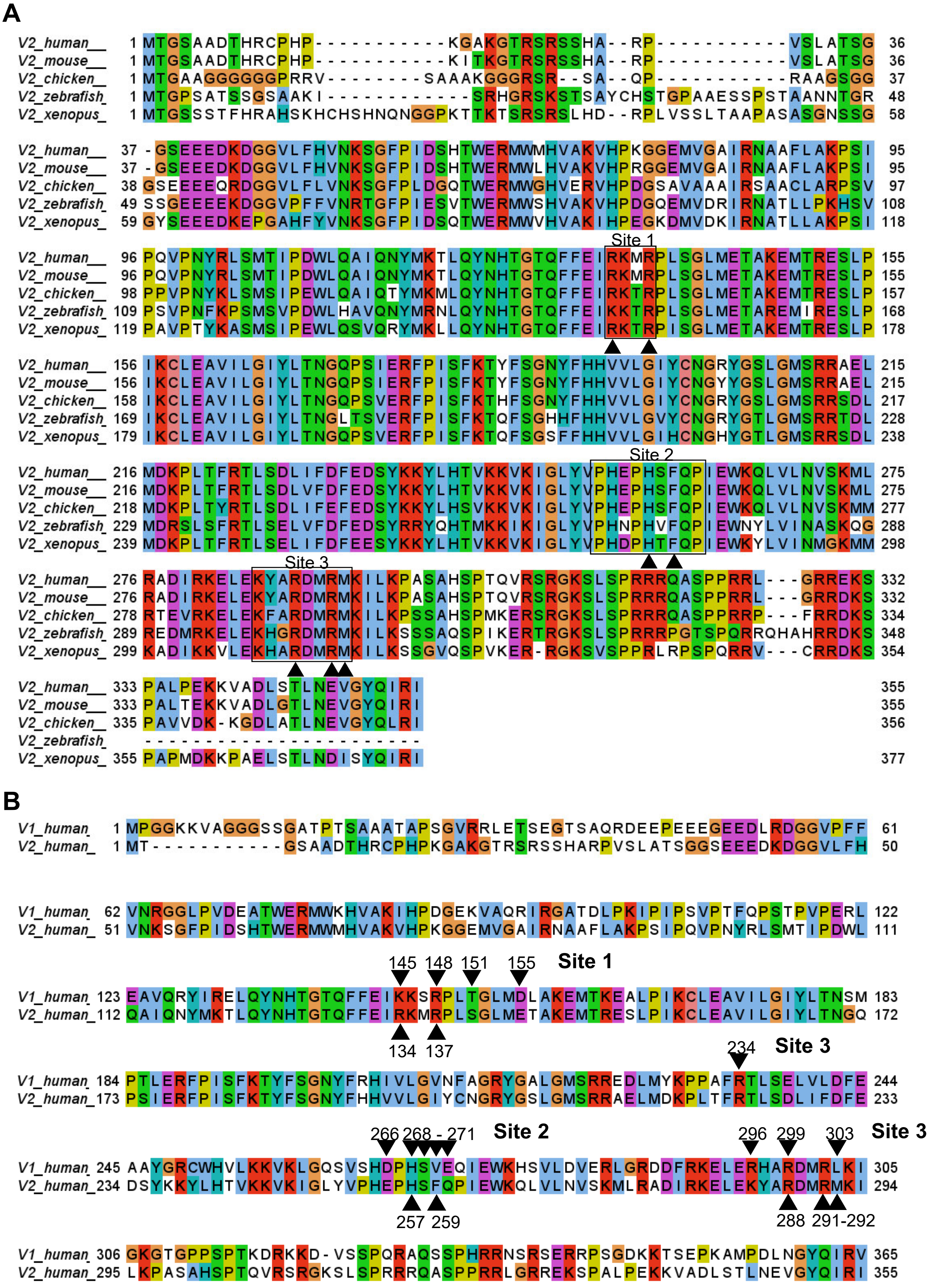
A
V2_human__ 1 MTGSAADTHRCPHP-----------KGAKGTRSRSSHA--RP-----------VSLATSG 36
V2_mouse__ 1 MTGSAADTHRCPHP-----------KITKGTRSRSSHA--RP-----------VSLATSG 36
V2_chicken_ 1 MTGAAGGGGGGPRRV--------SAAAKGGGRSR--SA--QP-----------RAAGSGG 37
V2_zebrafish 1 MTGPSATSSGSAAKI------------SRHGRSKSTSAYCHSTGPAEESSPSTAANNTGR 48
V2_xenopus_ 1 MTGSSSTFHRAHSKHCHSHNQNGGPKTTKTSRSRSLHD--RPLVSSLTAAPASASGNSSG 58
V2_human__ 37 -GSEEEDKDGGVLFHVNKSGFPIDSHTWERMWMHVAKVHPKGGEMVGAIRNAAFLAKPSI 95
V2_mouse__ 37 -GSEEEDKDGGVLFHVNKSGFPIDSHTWERMWLHVAKVHPRGGEMVGAIRNAAFLAKPSI 95
V2_chicken_ 38 GSEEEEQRDGGVLFLVNKSGFPLDGQTWERMWGHVERVHPDGSAVAAAIRSAACLARPSV 97
V2_zebrafish 49 SSGEEEEKDGGVPFFVNRTGFPIESVTWERMWSHVAKVHPDGQEMVDRIRNATLLPKHSV 108
V2_xenopus_ 59 GYSEEEDKEPGAHFYVNKSGFPIDSQTWERMWVHVAKIHPEGKDMVDKIRNATLLAKPSI 118
Site 1
V2_human__ 96 PQVPNYRLSMTIPDWLQAIQNYMKTLQYNHTGTQFFEIRKMRPLSGLMETAKEMTRESLP 155
V2_mouse__ 96 PQVPNYRLSMTIPDWLQAIQNYMKTLQYNHTGTQFFEIRKMRPLSGLMETAKEMTRESLP 155
V2_chicken_ 98 PPVPNYKLSMSIPEWLQAIQTYMKMLQYNHTGTQFFEIRKTRPLSGLMETAKEMTRESLP 157
V2_zebrafish 109 PSVPNFKPSMSVPDWLHAVQNYMRNLQYNHTGTQFFEIRKTRPLSGLMETAREMIRESLP 168
V2_xenopus_ 119 PAVPTYKASMSIPEWLQSVQRYMKLLQYNHTGTQFFEIRKTRPISGLMETAKEMTRESLP 178
V2_human__ 156 IKCLEAVILGIYLTNGQPSIERFPISFKTYFSGNYFHHVVLGIYCNGRYGSLGMSRRAEL 215
V2_mouse__ 156 IKCLEAVILGIYLTNGQPSIERFPISFKTYFSGNYFHHVVLGIYCNGYYGSLGMSRRAEL 215
V2_chicken_ 158 IKCLEAVILGIYLTNGQPSVERFPISFKTHFSGNYFHHVVLGIYCNGRYGSLGMSRRSDL 217
V2_zebrafish 169 IKCLEAVILGIYLTNGLTSVERFPISFKTQFSGHHFHHVVLGVYCNGRYGTLGMSRRTDL 228
V2_xenopus_ 179 IKCLEAVILGIYLTNGQPSVERFPISFKTQFSGSFFHHVVLGIHCNGHYGTLGMSRRSDL 238
Site 2
V2_human__ 216 MDKPLTFRTLSDLIFDFEDSYKKYLHTVKKVKIGLYVPHEPHSFQPIEWKQLVLNVSKML 275
V2_mouse__ 216 MDKPLTFRTLSDLVFDFEDSYKKYLHTVKKVKIGLYVPHEPHSFQPIEWKQLVLNVSKML 275
V2_chicken_ 218 MDKPLTYRTLSDLIFEFEDSYKKYLHSVKKVKIGLYVPHEPHSFQPIEWKQLVLNVSKMM 277
V2_zebrafish 229 MDRSLSFRTLSELVFDFEDSYRRYQHTMKKIKIGLYVPHNPHVFQPIEWNYLVINASKQG 288
V2_xenopus_ 239 MDKPLTFRTLSELIFDFEESYKKYLHTVKKVKIGLYVPHDPHTFQPIEWKYLVINMGKMM 298
Site 3
V2_human__ 276 RADIRKELEKYARDMRMKILKPASAHSPTQVRSRGKSLSPRRRQASPPRRL---GRREKS 332
V2_mouse__ 276 RADIRKELEKYARDMRMKILKPASAHSPTQVRSRGKSLSPRRRQASPPRRL---GRRDKS 332
V2_chicken_ 278 RTEVRKELEKFARDMRMKILKPSSAHSPMKERSRGKSLSPRRRQASPPRRP---FRRDKS 334
V2_zebrafish 289 REDMRKELEKHGRDMRMKILKSSSAQSPIKERTRGKSLSPRRRPGTSPQRRQHAHRRDKS 348
V2_xenopus_ 299 KADIKKVLEKHARDMRMKILKSSGVQSPVKER-RGKSVSPPRLRPSPQRRV---CRRDKS 354
V2_human__ 333 PALPEKKVADLSTLNEVGYQIRI 355
V2_mouse__ 333 PALTEKKVADLGTLNEVGYQIRI 355
V2_chicken_ 335 PAVVDK-KGDLATLNEVGYQLRI 356
V2_zebrafish -----------------------
V2_xenopus_ 355 PAPMDKKPAELSTLNDISYQIRI 377
B
V1_human 1 MPGGKKVAGGGSSGATPTSAAATAPSGVRRLETSEGTSAQRDEEPEEEGEEDLRDGGVPFF 61
V2_human_ 1 MT-----------GSAADTHRCPHPKGAKGTRSRSSHARPVSLATSGGSEEEDKDGGVLFH 50
V1_human 62 VNRGGLPVDEATWERMWKHVAKIHPDGEKVAQRIRGATDLPKIPIPSVPTFQPSTPVPERL 122
V2_human_ 51 VNKSGFPIDSHTWERMWMHVAKVHPKGGEMVGAIRNAAFLAKPSIPQVPNYRLSMTIPDWL 111
145 148 151 155
Site 1
V1_human 123 EAVQRYIRELQYNHTGTQFFEIKKSRPLTGLMDLAKEMTKEALPIKCLEAVILGIYLTNSM 183
V2_human_112 QAIQNYMKTLQYNHTGTQFFEIRKMRPLSGLMETAKEMTRESLPIKCLEAVILGIYLTNGQ 172
134 137
234
Site 3
V1_human 184 PTLERFPISFKTYFSGNYFRHIVLGVNFAGRYGALGMSRREDLMYKPPAFRTLSELVLDFE 244
V2_human_173 PSIERFPISFKTYFSGNYFHHVVLGIYCNGRYGSLGMSRRAELMDKPLTFRTLSDLIFDFE 233
266 268 - 271
Site 2
296 299 303
Site 3
V1_human 245 AAYGRCWHVLKKVKLGQSVSHDPHSVEQIEWKHSVLDVERLGRDDFRKELERHARDMRLKI 305
V2_human_234 DSYKKYLHTVKKVKIGLYVPHEPHSFQPIEWKQLVLNVSKMLRADIRKELEKYARDMRMKI 294
257 259
288 291-292
V1_human 306 GKGTGPPSPTKDRKKD-VSSPQRAQSSPHRRNSRSERRPSGDKKTSEPKAMPDLNGYQIRV 365
V2_human_295 LKPASAHSPTQVRSRGKSLSPRRRQASPPRRLGRREKSPALPEKKVADLSTLNEVGYQIRI 355

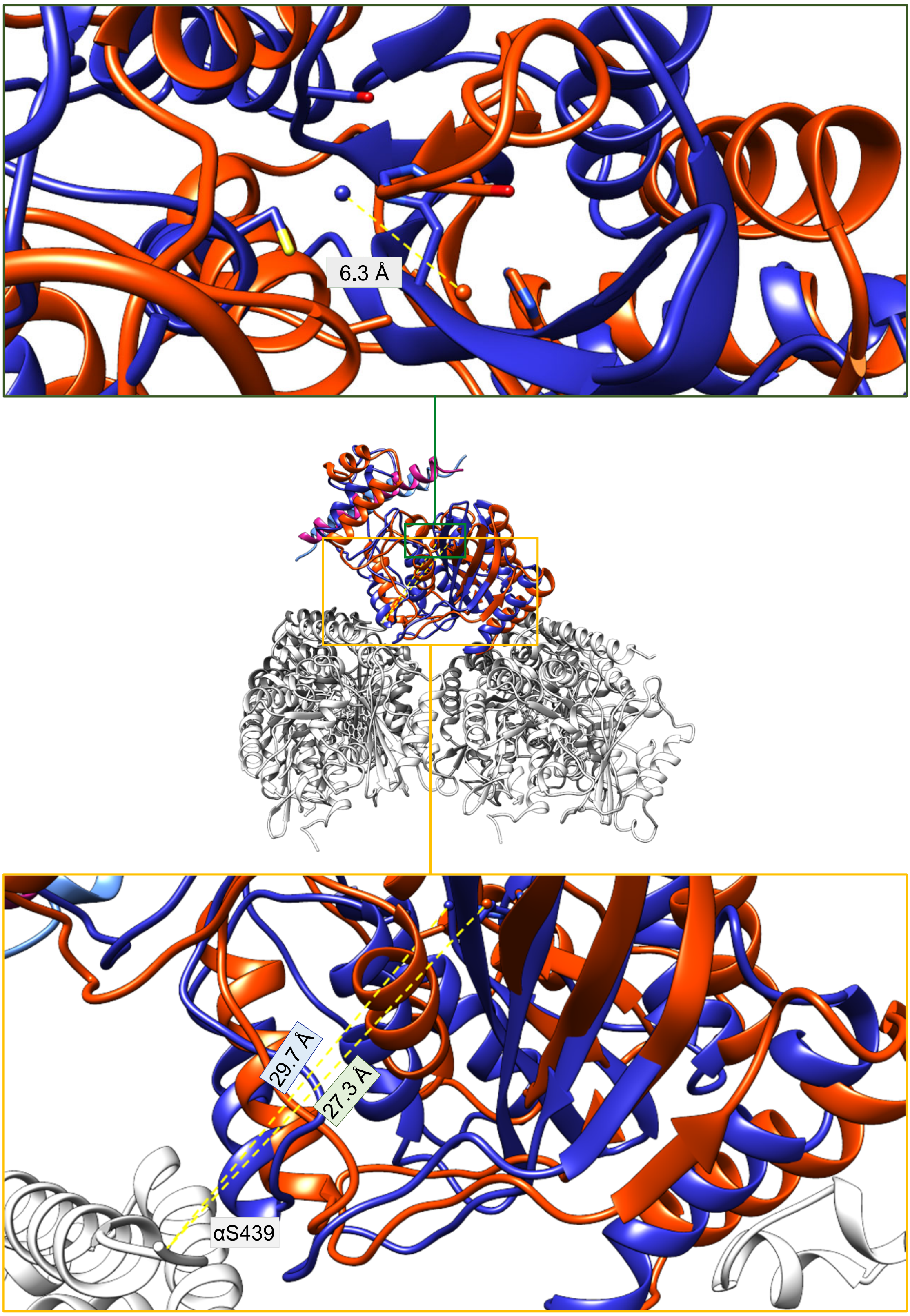
6.3 Å
29.7 Å
27.3 Å
αS439
